# Discovery and development of single-nucleotide polymorphism markers for resistance to *Striga gesnerioides* in cowpea (*Vigna unguiculata*)

**DOI:** 10.1101/2025.09.15.676209

**Authors:** Patrick Obia Ongom, Christian Fatokun, Ousmane Boukar

**Affiliations:** International Institute of Tropical Agriculture (IITA), Kano, Nigeria; International Institute of Tropical Agriculture (IITA), Ibadan, Nigeria

**Keywords:** Cowpea (*Vigna unguiculata*), *Striga gesnerioides*, genome-wide association study, candidate genes, SNP-based KASP markers, marker validation

## Abstract

The parasitic weed [*Striga gesnerioides* (Willd.) Vatke] is a principal biotic constraint to cowpea [*Vigna unguiculata* (L.) Walp.] production in West and Central Africa, causing severe yield reductions. Multiple races of *S. gesnerioides* exist across the cowpea-growing areas of the sub- region. Past efforts identified some resistant sources and race-specific genes underpinning Striga resistance, but deployment of associated markers in breeding is limited. Here, we utilized a 51K cowpea iSelect single-nucleotide polymorphisms (SNPs) to decipher genomic regions underlying Striga resistance and explore marker conversion and validation for easy deployment. The study used two-year phenotypic data on a minicore panel of 368 cowpea genotypes screened at two sites in Northern Nigeria. SNPs performances were verified and validated using two independent sets of 60 and 20 diverse genotypes respectively. The minicore displayed apparent differences in response to the *S. gesnerioides* attack. A genome-wide scan uncovered a primary gene effect signal on chromosome Vu11 and minor regions on chromosomes Vu02, Vu03, Vu07, Vu09 and Vu10. The major effect region on Vu11 harbored a coil-coil nucleotide-binding site leucine-rich repeat (CC- NBS-LRR) protein, encoded by the *RSG3-301* gene, previously implicated in race-specific resistance to *S. gesnerioides* in cowpea. The associated SNPs were successfully converted into Kompetitive Allele-Specific PCR (KASP) assays and validated using 20 independent diverse cowpea genotypes. Five KASP markers, snpVU00075, snpVU00076, snpVU00077, snpVU00078, and snpVU00079, depicted consistent and significant associations with the phenotype in the validation set. The markers provide valuable tools for efficient marker-assisted selection (MAS) in breeding programs focused on developing Striga-resistant cowpea varieties.

## 1 Introduction

Cowpea (*Vigna unguiculata* L. Walp.) is a vital legume crop widely grown in sub-Saharan Africa, Asia, and Latin America (Abate et al., 2012; Singh et al., 2014). It is a self-pollinated diploid with 2n=22 chromosomes and a genome size of 640.6 Mbp (Lonardi et al., 2019). Cowpea has synonymous names worldwide, including black-eyed pea, lubia, Kathir pea, China pea, crowder pea, niebe, and southern pea (Heuzé et al., 2015). Recent evidence traces the cowpea center of domestication to Nigeria in West Africa (WA) (Vaillancourt and Weeden, 1992; Panzeri et al., 2022). The crop is grown in over 88 countries worldwide, with Nigeria being the largest producer, accounting for 46% of world production. Cowpea is an essential food and income source for more than 200 million smallholder farmers, particularly in semi-arid regions where other crops may not thrive due to limited water availability (Abate et al., 2012; Singh, 2014; Silva et al., 2018). Cowpea is a highly nutritious crop rich in protein, dietary fiber, vitamins, and minerals (Singh et al., 2014; Jayathilake et al., 2018). The crop is also an essential source of nitrogen for the soil, making it a valuable component of sustainable cropping systems (Blade et al., 1997; Mortimore et al., 1997); it serves as feed for livestock and as a cover crop to protect and enrich the soil (Abate et al., 2012).

Despite its importance, cowpea production is often constrained by a range of biotic and abiotic stresses, including parasitic weeds (Striga and Alectra), insect pests and diseases, drought, and poor soil fertility (Atokple et al., 1995; Fatokun et al., 2002; Li et al., 2009; Boukar et al., 2013; Nkomo et al., 2021). Among these constraints, the parasitic weed [*Striga gesnerioides* (Willd.) Vatke], is quite devastating on cowpea, especially in the Sudano-Sahelian belt of West Africa, where the crop is most widely cultivated (Parker and Riches, 1993; Mohamed et al., 2001; Li et al., 2009; CABI, 2020). Cowpea yield loss to *S. gesnerioides* ranges from moderate to total crop loss in some parts of Nigeria, Niger, and Burkina Faso (CABI, 2020).

*Striga gesnerioides* belongs to the Orobanchaceae family (CABI, 2020). It is an obligate parasite with tiny seeds, unable to establish itself without the help of a host plant CABI, 2020). Germination depends on a period of moist conditioning and exposure to germination stimulants in the host plant’s root exudates, the most important of which is alectrol, a stimulant for both the Striga and related parasite *Alectra vogelii* (Müller et al., 1992; CABI, 2020). Over 28 Striga species and six subspecies have been characterized, among which purple witchweed [*Striga hermonthica* (Delile) Benth.], Asiatic witchweed [*Striga asiatica* (L.) Kuntze], and *S. gesnerioides* are the most economically important (Mohamed et al., 2001). *Striga hermonthica* and *S. asiatica* primarily infect cereals in the *Poaceae* family, while the primary hosts of *S. gesnerioides* are cowpea and wild legume species (Ohlson and Timko, 2020). However, *S. gesnerioides* can parasitize hosts across genera, including *Ipomea, Jaquemontia, Merremia, Euphorbia, and Nicotiana* (Mohamed et al., 2001).

Several control strategies have been developed for parasitic weeds, including improved cultural practices, breeding using wild and cultivated germplasm as sources of resistance, and the use of chemical control, but the use of resistant cultivars is still considered the most effective (Parker and Riches, 1993; Singh et al., 2006; Li et al., 2009). Given its significance in cowpea production*, S. gesnerioides* resistance is required for varieties released for WA’s Sahelian and Sudan Savanna zones. Hence, Striga resistance is a “must-have trait” in the cowpea product profiles. Consequently, significant efforts have been made to study the Striga race structure and identify resistance sources in cowpea. Molecular profiling and host differential response studies using *S. gesnerioides* isolates across the WA region revealed seven distinct parasite races (Botanga and Timko, 2006). An additional report by (Li et al., 2009; Ohlson and Timko, 2020) confirmed the seven races of *S. gesnerioides*. The authors observed that some cowpea cultivars were differentially resistant to various geographic isolates of the parasite. The races have been designated as SG1 (Burkina Faso), SG2 (Mali), SG3 (Nigeria and Niger), SG4 (Benin), SG4z (localized to the Zakpota region of Benin), SG5 (Cameroon), and SG6 (Senegal) (Li et al., 2009). However, Ohlson and Timko, (2020) reported that SG6 from Senegal was related to the SG1 race, and the authors revealed a novel race (designated as SG6) in Nigeria that was able to overcome more cowpea resistance genes than any previously reported race.

Screening efforts have identified sources of resistance to *S. gesnerioides*. These include B301 (Botswana landrace), IT82D-849, IT81D-994, and Wango-1 (from Burkina Faso), all having monogenic dominant inheritance mode and are resistant to races SG1, SG2, and SG3 (Singh and Emechebe, 1990; Atokple et al., 1993; Li et al., 2009). Genotype HTR (from Niger) was reported to carry one or two dominant genes and is resistant to race SG1, while Suvita-2 has a single recessive gene against SG3 and a single dominant gene against SG1 and SG2 (Li et al., 2009). In addition, Omoigui et al. (2010) screened some cowpea genotypes and found B301, IT03K-338-1, and IT99K- 573-2-1 to be free of emerged Striga and Alectra shoots, while IT98K-1092-1 and IT97K-205-8 were resistant to Striga but supported the emergence of some Alectra shoots.

Past efforts also identified some molecular markers associated with resistance to *S. gesnerioides* in cowpea. Notably, Ouédraogo et al. (2001), (2002); Li et al. (2009) reported amplified fragment length polymorphism (AFLP) markers on linkage group 1 (LG1) linked to race-specific genes: *Rsg2- 1* in the cowpea line IT82D-849, *Rsg1-1* in B301, and *Rsg4-3* in Tvu14676. Additionally, race-specific genes *Rsg3-1* and *Rsg994-1*, present in Suvita-2 and IT91D-994 respectively, were mapped on LG6 (Ouédraogo et al., 2001; Li et al., 2009). Two sequence-characterized amplified regions (SCARs) markers 61R (E-ACT/M-CAA) and SEACTMCAC83/8, linked to *S. gesnerioides* resistance, were developed and deployed for marker-assisted selection (Ouédraogo et al., 2001; Li et al., 2009; Ouédraogo et al., 2012). Four AFLP markers, E-ACT/M-CTC_115_, E-ACT/M-CAC_115_, E- ACA/M-CAG_108,_ and E-AAG/E-CTA_190_, were found to be associated with the *Rsg1* gene in a resistant line IT93K- 693-2 that confers resistance to SG3 race (Boukar et al., 2004). Some Simple Sequence Repeat (SSR) markers associated with resistance to *S. gesnerioides* race 3 (SG3) were identified and deployed in breeding for resistance (Li et al., 2009; Asare et al., 2013; Omoigui et al., 2017; Essem et al., 2019). However, these old marker technologies have known limitations for wide-scale routine applications in breeding, including difficulty in handling and scoring, and a lack of automation to allow high-throughput genotyping. Consequently, there has been a significant shift toward using SNPs, given their genomic abundance and rapid emergence of novel, faster, and cheaper methods of genotyping (Tsuchihashi and Dracopoli, 2002).

Recent progress in developing next-generation genomics and genetic resources for cowpea has generated more robust molecular marker platforms than those described earlier. Notably, Muchero et al. (2009) developed a cowpea GoldenGate assay consisting of 1536 SNP markers, being utilized for linkage mapping and QTL analyses (Lucas et al., 2011; Muchero et al., 2013; Pottorff et al., 2014) and assessment of genetic diversity (Huynh et al., 2013). Illumina Cowpea iSelect Consortium Array having 51,128 SNPs, was also developed (Muñoz-Amatriaín et al., 2017) and has been deployed in genome-wide mapping of several traits in cowpea (Herniter et al., 2018, 2019; Lo et al., 2018, 2019; Miesho et al., 2019; Paudel et al., 2021; Ongom et al., 2022b). Recently, cowpea researchers have developed low-density Kompetitive Allele Specific PCR (KASP) assays (Ongom et al., 2021; Wu et al., 2021) and medium-density DArTag genotyping panel (Ongom et al., 2022a, 2024). Several cowpea genetic resources have also been developed; among them are the UCR minicore, a diverse set of genotypes that have been genotyped with the iSelect SNPs panel (Muñoz-Amatriaín et al., 2021), and the IITA minicore, a set of genotypes representing the cowpea germplasm maintained at the IITA Genetic Resources Center that have been genotyped based on genotyping by sequencing (GBS) (Fatokun et al., 2018). The cowpea genomic and genetic resources so far developed have opened doors for QTL/gene discovery and development of robust molecular markers to enhance breeding for essential traits, including Striga resistance.

Resistance to *Striga gesnerioides* is notoriously difficult to evaluate in the field due to a complex interplay of confounding factors: seasonal shifts, soil heterogeneity, subjective scoring criteria, and significant parasite race diversity (Li et al., 2009). These challenges highlight a critical need for a reliable marker-assisted selection (MAS) platform, especially given the prevalence of multiple *S. gesnerioides* races that may circumvent resistance derived from a single source. In response, this study aimed to: (i) identify SNPs robustly associated with Striga resistance; and (ii) develop Kompetitive Allele Specific PCR (KASP) assays suitable for routine use in cowpea breeding. The study harnessed the diversity in the UCR minicore panel, combining two years of Striga resistance phenotypic data with the high-density iSelect SNPs to pinpoint genomic regions associated with *S. gesnerioides* in cowpea. Ultimately, the development and validation of KASP markers linked to Striga resistance will advance the integration of MAS into breeding pipelines. These markers offer breeders a precise, cost-effective tool for selecting Striga resistance, expediting the development of resilient cowpea varieties and accelerating their deployment in farmers’ fields.

## 2 Materials and methods

### 2.1 Genetic materials

The study used 368 UCR minicore genotypes described by (Muñoz-Amatriaín et al., 2021). The UCR minicore contains diverse landraces and breeding materials from 50 countries covering Africa, Asia, North and South America, Europe, and Australia. The minicore population, previously genotyped with high-density SNPs, represents the existing genetic and phenotypic diversity of cultivated cowpea while maintaining a sample size that can be managed by most researchers and breeders for evaluating traits of interest (Muñoz-Amatriaín et al., 2021).. For marker development, the present study used 60 diverse cowpea genotypes along with 46 F_1_ progenies for SNPs technical verification and another independent set of 20 genotypes consisting of some known resistant and susceptible sources commonly used as standard checks in breeding programs for marker validation.

### 2.2 Study sites

The study was conducted at three sites in Northern Nigeria where there is a widespread of *Striga gesnerioides*. The first site was a Striga hot spot at IITA Minjibir Research Farm, Kano, Latitude 12° 14′ 35.30″ N and Longitude 8° 66′ 62.10″ E. The second Striga hot spot site was at Malam Madori, Jigawa, located at Latitude 12° 33’ 36.32” N and Longitude 9° 59’ 9.56” E. Previously, the samples of Striga races collected from these two sites have been characterized as belonging to race *SG3*, which is a dominant *S. gesnerioides* race in Nigeria among other less frequent races labeled as *SG1, SG5* and *SG6* (Ohlson and Timko, 2020). The third site designated for validation screening is situated at the IITA station in Kano, located at latitude 11°58’50.0"N and longitude 8°33’26.8"E. This facility includes a nursery that has been deliberately infested with *Striga gesnerioides* race SG3 to facilitate controlled research on cowpea resistance to this parasitic weed.

### 2.3 Striga resistance phenotyping

The minicore genotypes were evaluated in the two locations in Northern Nigeria for two years in Minjibir and one year in Malam Madori. The genotypes were planted in a two-row plot of 4 meters long and at a spacing of 0.75 m between the rows and 0.2 m within the row. The trials were established as alpha lattice designs with two replications. Variation in Striga infestation was assessed using two criteria: First, the Striga score was evaluated on a scale of 0-3, where 0 = no Striga emergence; 1 = few Striga emerged; 2 = moderate Striga emergence; 3 = heavy Striga infestation The second method was the presence-absence rating, where ‘0’ indicates that Striga is absent in the plot and ‘1’ indicates Striga is present. In addition, a validation set of 20 cowpea genotypes was screened in an artificially infested nursery using a randomized complete block design experiment. For the past several years, this nursery has been dedicated to Striga screening following heavy inoculation of the soil with seeds of Striga collected from infested fields at Malam Madori, Jigawa State, Nigeria. Prior to soil inoculation, Striga seeds were preconditioned through surface-sterilization using 10% sodium hypochlorite for ∼10 minutes followed by incubation in the dark at 29 °C for 10 days. The soil was then inoculated with Striga seeds at a rate of 2 g/m^2^, evenly distributing the seeds across the soil surface. The 20 cowpea genotypes were each planted in a 2-row plot of 4 m and spaced at 0.75 m by 0.2 m between and within rows, respectively, and the experiment had two replications. The presence or absence of Striga was assessed for each plot, including the number of Striga plants that emerged, and the data was analyzed with Analysis of Variance (ANOVA) and means compared using LSD test.

### 2.4 SNP genotype data

The SNP data was obtained from (Muñoz-Amatriaín et al., 2017). The genotyping was done at the University of Southern California Molecular Genomics Core lab (Los Angeles, California, USA) following the procedure described by (Muñoz-Amatriaín et al., 2021). Briefly, genomic DNA was extracted from the young leaves of individual plants using DNeasy Plant Kit (Qiagen, Valencia, California, USA). The Cowpea iSelect Consortium Array including 51,128 SNPs (Muñoz-Amatriaín et al., 2017) was used for genotyping each DNA sample. SNPs were called in GenomeStudio (Illumina Inc., San Diego, California, USA) and manually curated to remove those with high levels (>20%) of missing data and/or heterozygous calls. The genotype data contained 47129 SNPs after removing the contigs with no chromosomal assignment. Further filtering was conducted, setting the minimum minor allele frequency at 0.05 and maximum heterozygous proportion at 0.1, resulting in 41510 SNPs for downstream analyses.

### 2.5 Data analysis

The phenotypic data were analyzed using a general linear model tailored for an alpha lattice design, implemented through the agricolae package in R (Mendiburu, 2020). The model applied is represented as:

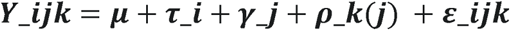

Where: Y__jk denotes the observed phenotypic value for the *i*^th^ treatment in the *k*^th^ block within the *j*^th^ replication., µ is the general mean, __ represents the fixed effect of the *i*^th^ treatment, _j signifies the effect of the *j*^th^ replication, and p_kkj) accounts for the effect of the *k*^th^ incomplete block nested within the *j*^th^ replication, __jk is the random error term associated with the observation. The mean values for each minicore accession were extracted from the model using the LSD.test () function and later utilized in GWAS analysis.

GWAS was conducted using the 41510 filtered SNPs in TASSEL v 5.2.20 (Bradbury et al., 2007) and rMVP package v 1.03 (Yin et al., 2021), utilizing the Mixed Linear Model (MLM) (Zhang et al., 2010) and the Fixed and random model Circulating Probability Unification (FarmCPU) (Liu et al., 2016) on the following traits: Striga score and Striga presence-absence rating. The FarmCPU model uses a multiple loci linear mixed model (MLMM) and incorporates multiple markers simultaneously as covariates in a stepwise MLM to partially remove the confounding between testing markers and kinship (Liu et al., 2016). A genomic PCA matrix (P) and kinship matrix (K) were used to capture the population structure and relatedness among individuals in the panel (Kang et al., 2008). In TASSEL, kinship, and principal components were computed and fitted together with the SNPs in the MLM model to account for relatedness and population structure, respectively. The MLM statistics were also utilized separately in CMplot package 3.1.3 (Yin, 2016) to generate customized GWAS Manhattan and QQ plots. Decisions on significant GWAS signals were based on both the conservative Bonferroni (Henry, 2015) and a less conservative false discovery rate (FDR) (Glickman et al., 2014; Ongom et al., 2022b, 2024) corrections of multiple statistical tests to reduce the risk of a type I error.

### 2.6 Candidate gene search

Candidate genes linked to Striga resistance were pinpointed by aligning the positions of significant SNPs from the GWAS with the cowpea reference genome (version 1.1) available on Phytozome 13 Genome Browser (https://phytozome-next.jgi.doe.gov/info/Vunguiculata_v1_1, accessed on 26 January 2025). A length of 200 kb was added or removed from either end of the significant marker to locate potential regions for comparison based on the LD rate of the cowpea minicore population (Ongom et al., 2022b). In selecting candidate genes, the following criteria were used: (i) genes of known function in cowpea related to the trait under study, (ii) genes with function-known orthologs in Arabidopsis related to the trait under study, and (iii) genes pinpointed by the peak SNPs. Putative candidate genes were subsequently researched in the literature for verification. We then used chromoMap v4.1.1 package (Anand and Rodriguez Lopez, 2022) to visualize the distribution of selected genes on target chromosomes where GWAS signals were discovered.

### 2.7 KASP marker development

Technical verification of KASP assay: Seventeen SNP markers spanning Striga resistance gene regions were selected to develop KASP assays. The design sequences of the selected SNPs are presented in **Table 1**. The validity of these KASP assays was verified using 60 cowpea genotypes and 46 heterozygous F_1_ progenies. The samples for this SNP technical verification were collected in three replicates per genotype. The leaf samples and SNP design sequences were sent to Intertek Lab Sweden for assay development. Leaf sampling followed the standard procedure previously described (Ongom et al., 2021). Genotyping data were visualized using SNPviewer version 5.1.1.27582 (LGC Genomics, 2025) and SNP cluster plots were generated.

**Table 1.**
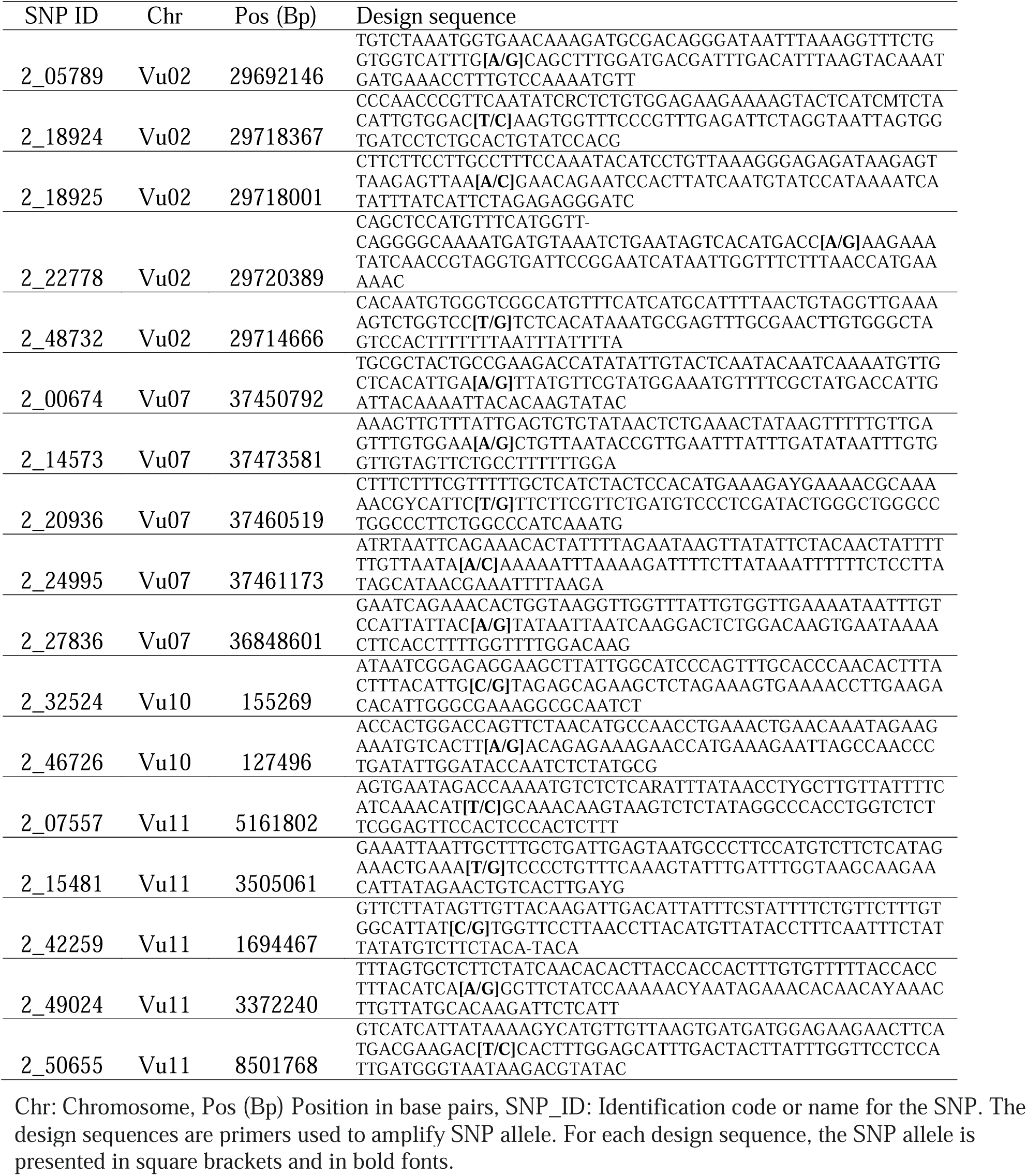
Candidate SNP markers for Striga resistance used to develop the KASP assay.

Testing the marker-trait association: To validate the performance of these markers for tracking Striga resistance, we deployed the candidate markers in a panel of 20 cowpea validation genotypes consisting of some known resistant and susceptible sources. These cowpea genotypes were independently phenotyped in a an artificially Striga-infested nursery to confirm their reaction status. The number of Striga plants that emerged in each plot was counted, and the data was used to categorize the 20 genotypes into either resistant (R) or susceptible (S) with zero Striga count = R and any count greater than zero = S. The 20 cowpea genotypes were then genotyped with the 17 candidate KASP markers. A single marker analysis was then conducted to verify the consistency of these candidate markers in tracking Striga resistance in cowpea. A chi-square test of independence and a t-test were performed in R for each marker with the null hypothesis that each marker genotype is independent of the observed variation in the phenotypic status of the cowpea genotypes. The results of these tests were visualized using plots generated by ggplot2 and ggstatsplot packages.

## 3 Results

### 3.1 Phenotypic assessment

#### Phenotypic assessment

The phenotypic data from both Minjibir and Malam Madori sites depicted skewed distributions for Striga ratings, with a greater proportion of the minicore genotypes showing susceptibility to Striga. This is portrayed by the stacked bar chart in **Figure 1A**, where only 2.7 to 6.5% of the minicore showed resistance across the two testing sites.. In contrast, a mean Striga score of 2.35 (scale 1-3) was registered for the combined data across the two locations **(Figure 1B)**. The susceptible genotypes triggered extensive growth of Striga plants, causing severe yellowing and death of cowpea plants. At the same time, no emergence of Striga was observed in plots planted with resistant genotypes **(Figure 1C).**

**Figure 1.**
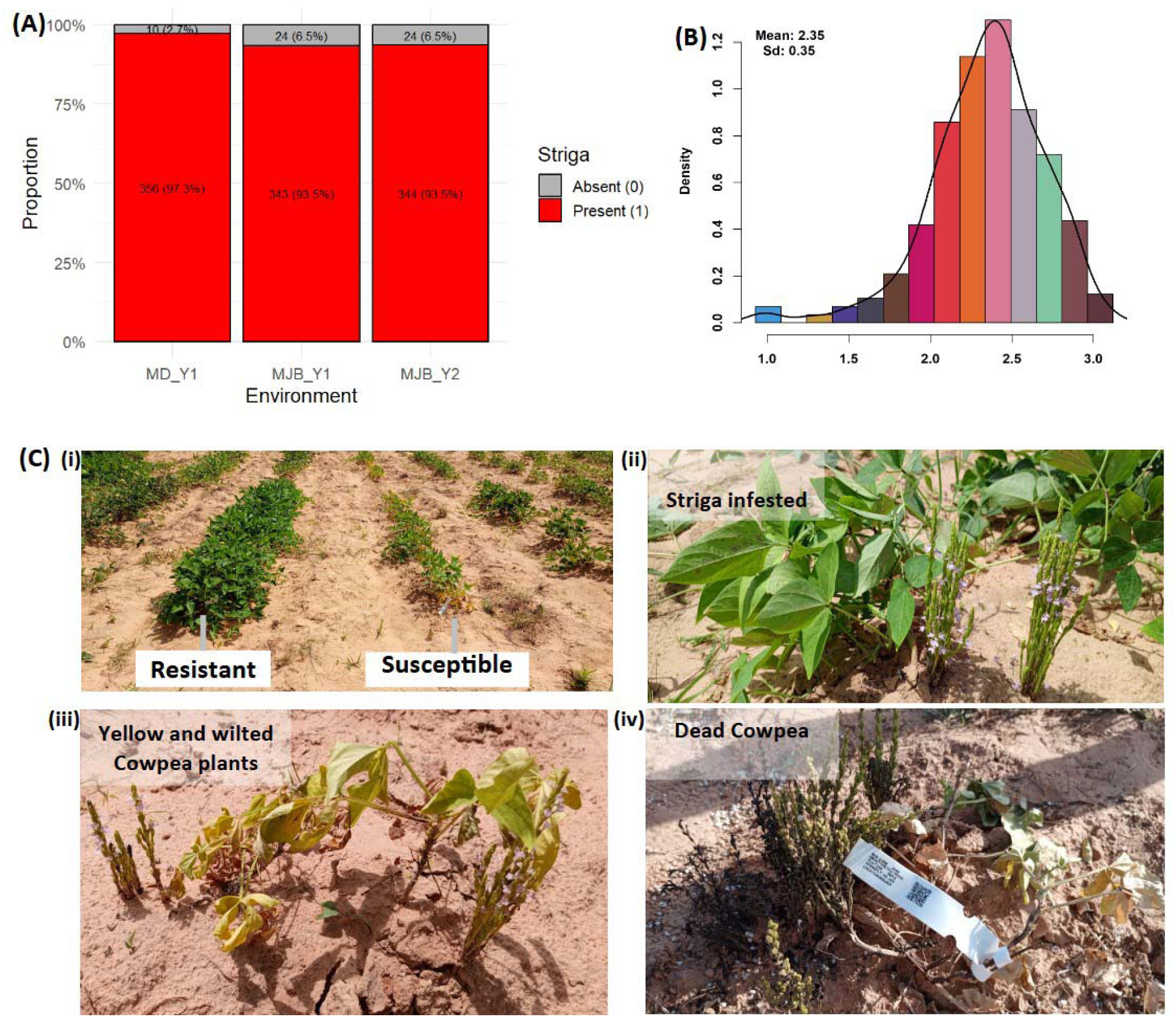
Density curve displaying the phenotypic distribution and reaction of minicore genotypes to Striga gesnerioides. **(A)** the distribution for presence-absence rating at Minjibir with the x-axis indicating a Striga rating of 0 (Striga absent) and 1 (Striga present), **(B)** the distribution for presence- absence rating at Malam Madori with the x-axis indicating Striga rating of 0 (Striga absent) and 1 (Striga present) (**C)** the distribution for Striga score (on the x-axis) for combined data across the two locations, (**D)** the response of minicore genotypes to Striga infestation in the field, with panel (i) depicting resistant and susceptible genotypes and panels (ii) to (iv) displaying the phases of plant growth under Striga infestation from green healthy-looking but infested plants through severe yellowing and wilting up to completely dead cowpea plants.

### 3.2 Association analysis

GWAS discovered one major association signal for Striga resistance on chromosome Vu11 that was consistently significant in both locations and five other minor signals on chromosomes Vu02, Vu03, Vu07, Vu09, and Vu10 (**Figure 2A**). The major association signal on chromosome Vu11 and two minor regions on chromosomes Vu07 and Vu10 were detectable in both Minjibir and Malam Madori locations, while the regions on Vu03, Vu02, and Vu09 were significant only in one location (**Figure 2A**). The QQ plot shows that the observed p-values largely align with the expected distribution under the null hypothesis, indicating no evidence of inflation. However, the upward deflection at the tail suggests the presence of a small number of SNPs exhibiting stronger association signals than expected by chance.” (**Figure 2B**). It was also evident from the QQ plot that the GWAS signals were stronger in Malam Madori than in Minjibir and the combined location data set for Striga scores (**Figure 2B**).

**Figure 2.**
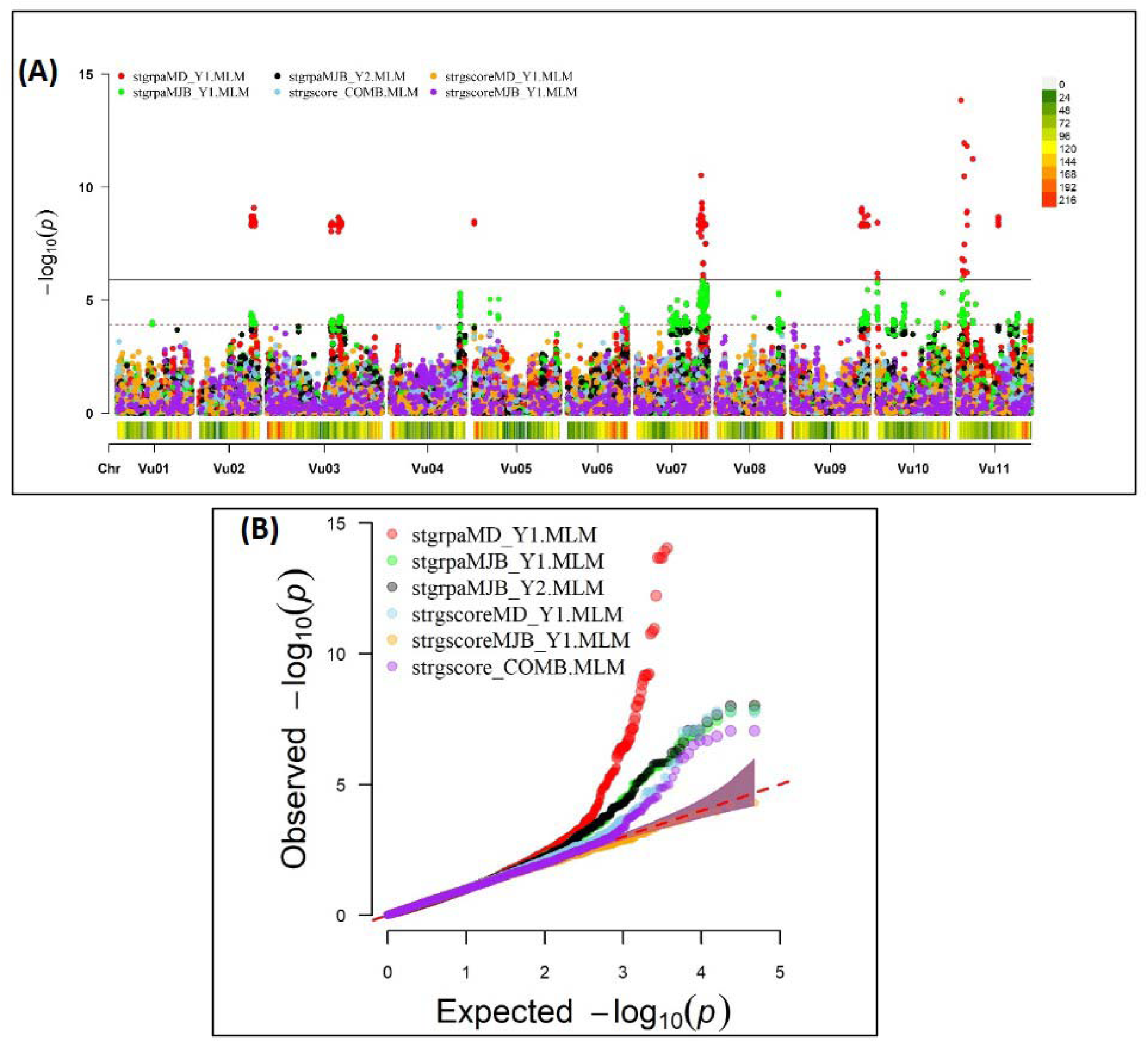
Genome-wide association signals based on the mixed linear model (MLM). **(A)** Manhattan plot depicting a major GWAS signal on chromosome Vu11 and five minor ones on chromosomes Vu02, Vu03, Vu07, Vu09 and Vu10. The data is presented for Striga presence-absence rating in Minjibir (stgrpaMJB_Y1.MLM and stgrpaMJB_Y2.MLM) and Malam Madori (stgrpaMD_Y1.MLM), Striga score in Malam Madori (stgrscoreMD_Y1), Striga score in Minjibir (stgrscoreMJB_Y1.MLM) and Striga score for data combined across the two locations (atgrscore_COMB.MLM). Beneath each chromosome in the Manhattan plot is the SNP density and distribution defined by the coloured key on the top right, with gray color representing low SNP density while red depicts high SNP density regions. (**B)** the QQ plot for the six experimental data sets.

A total of 309 significant SNPs with the Boneforrini threshold of -log10(p) > 5.92 were found to be associated with Striga resistance (**Supplementary Table 1**). Out of these, seventeen (17) representative SNPs that had R^2^ values above 7.5% were selected for KASP assay development (**Table 2**). The strongest GWAS signal on chromosome Vu11 was represented by five significant SNPs, accounting for 15.8% to 19.8% of phenotypic variance (**Table 2**). The other minor signals on chromosomes Vu02, Vu03, Vu07, Vu09, and Vu10 explained 6.8% to 14.1% of phenotypic variance (**Supplementary Table 1**).

**Table 2.** Representative SNPs that were significantly associated with Striga resistance in the cowpea minicore population.

| SNP Marker | Chr | Pos (bp) | R<sup>2</sup> | -log10(p) |
| --- | --- | --- | --- | --- |
| 2_05789 | Vu02 | 29,692,146 | 8.0% | 6.03 |
| 2_18924 | Vu02 | 29,718,367 | 7.9% | 6.02 |
| 2_18925 | Vu02 | 29,718,001 | 7.9% | 6.02 |
| 2_22778 | Vu02 | 29,720,389 | 8.0% | 6.03 |
| 2_48732 | Vu02 | 29,714,666 | 7.9% | 6.02 |
| 2_00674 | Vu07 | 37,450,792 | 11.4% | 8.49 |
| 2_14573 | Vu07 | 37,473,581 | 11.4% | 8.49 |
| 2_20936 | Vu07 | 37,460,519 | 11.4% | 8.48 |
| 2_24995 | Vu07 | 37,461,173 | 11.4% | 8.49 |
| 2_27836 | Vu07 | 36,848,601 | 12.8% | 9.50 |
| 2_32524 | Vu10 | 155,269 | 14.1% | 11.30 |
| 2_46726 | Vu10 | 127,496 | 10.4% | 8.67 |
| 2_07557 | Vu11 | 5,161,802 | 19.0% | 13.69 |
| 2_15481 | Vu11 | 3,505,061 | 18.0% | 14.03 |
| 2_42259 | Vu11 | 1,694,467 | 19.8% | 14.19 |
| 2_49024 | Vu11 | 3,372,240 | 15.8% | 12.44 |
| 2_50655 | Vu11 | 8,501,768 | 17.0% | 12.34 |

### 3.3 Candidate genes

Searching candidate genes within 200kb of peak GWAS signals identified 178 genes on four target chromosomes, with 28 on Vu11, 38 on Vu10, 56 on Vu07, and 57 on Vu02 **(Supplementary Table 2)**. Sixty-four of the 178 genes were clustered closely to the peak SNPs on the target chromosomes **(Figure 3)**. Further examinations revealed 20 unique annotated proteins associated with the 64 genes (**Figure 3),** given that most of these gene loci encoded similar functional proteins. Of the 64 candidate genes identified, 11 were positioned within approximately -167461 bp to 191285 bp of the lead SNP on Chromosome Vu11, five genes were within ∼ -139927 bp to 83997 bp of the association peak on Vu10, 36 genes were within ∼ -190061 bp to 202188 bp of the signal on Vu07 and 12 were within ∼ -145716 bp to 125678 bp of the association locus on Vu02 **(Table 3).** Several of these clusters contain genes with related biological functions.. For instance, up to five genes on Chromosome Vu02 belonged to the Pentatricopeptide repeat (PPR) superfamily protein. On chromosome Vu07, up to 28 genes had functions related to Cysteine-rich receptor-kinase-like protein, while 10 genes on chromosome Vu11 had functions related to LEUCINE-RICH REPEAT-CONTAINING PROTEIN **(Table 3).** The rest of the genes on each chromosome had unique functions, including, among others, C2H2-like zinc finger protein, myb transcription factor, MADS-box transcription factor family protein, ETHYLENE-RESPONSIVE TRANSCRIPTION FACTOR ERF003, and WRKY DNA -binding domain (WRKY) **(Table 3**, **Figure 3)**.

**Figure 3.**
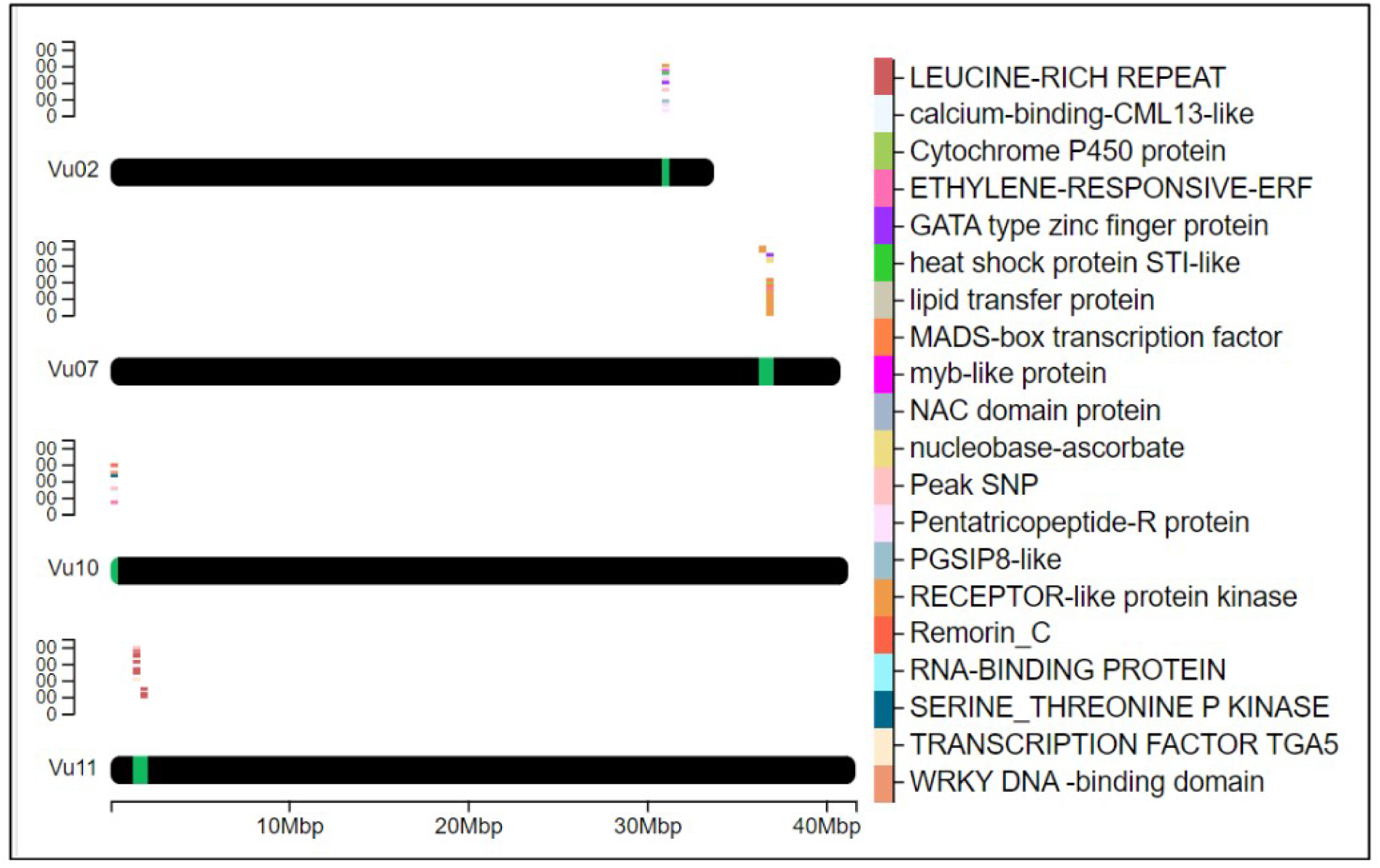
Heatmap of the position of key genes on chromosomes. It depicts the selected proximal genes to the peak SNPs with likely functional involvement in plant defense and immune signaling on chromosomes Vu02, Vu07, Vu10, and Vu11. The chromosomal positions of the genes are marked in green, while a cluster plot of genes is displayed above the marked regions. The color-coded legend on the right identifies 20 unique proteins associated with genes found within the QTL signal region.

**Table 3.**
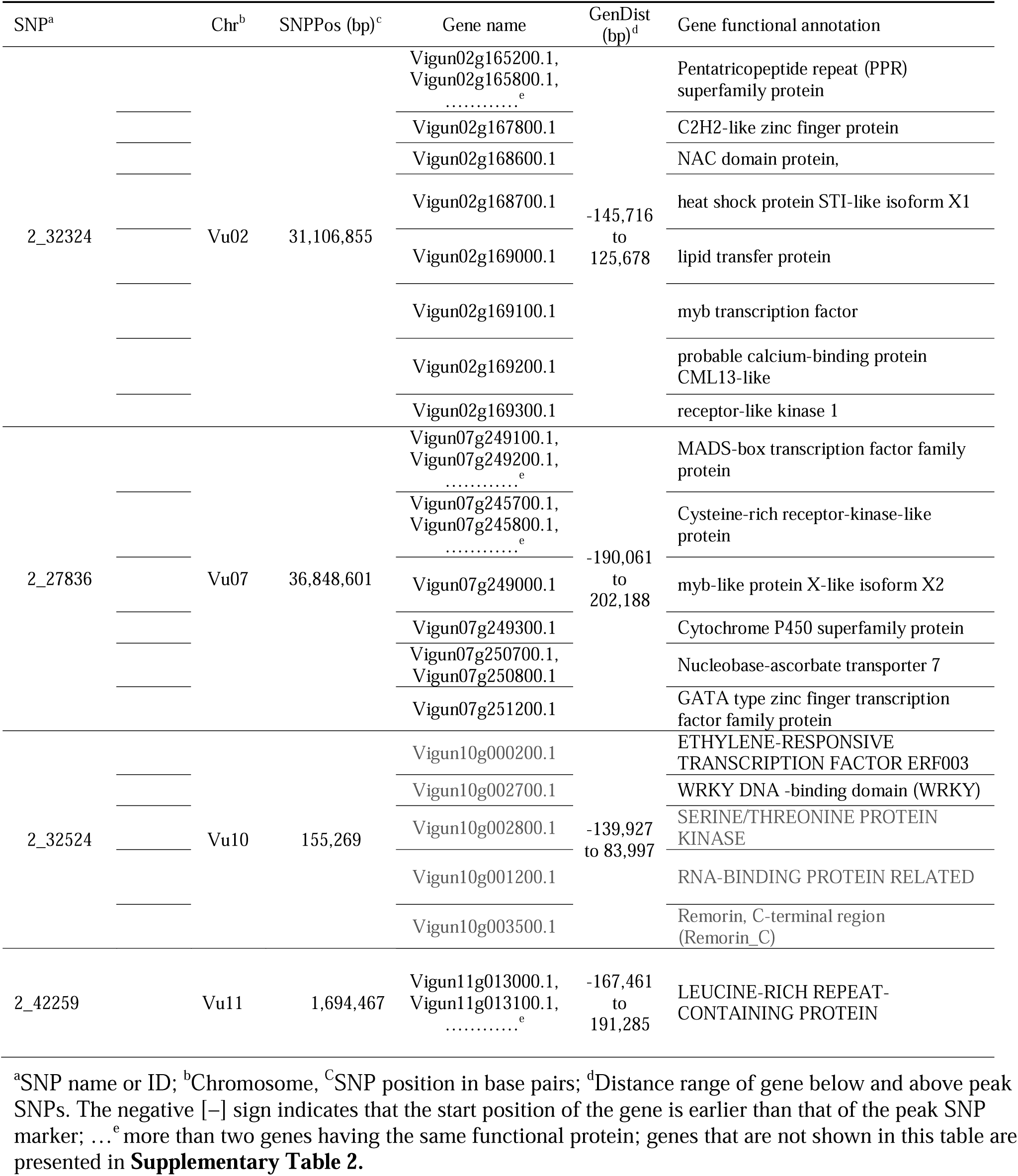
Selected genes found within 200kb of the representative GWAS SNPs with implicated functions in plant immunity and defense signaling.

### 3.4 KASP marker assay development

KASP assays were successfully developed for the 17 representative SNPs having strong associations with Striga resistance in cowpea. The SNP calls were verified using a unique technical validation set of cowpea genotypes, including homozygous genotypes and highly heterozygous F_1_. The raw genotype calls displaying the KASP assay performance of the 17 candidate SNPs in the cowpea technical validation set are provided in **Supplementary Table 3**. The KASP assay technical verification results revealed 12 SNPs as having good quality assays, allowing easy scoring of the alleles **(Figure 4A)**. Two of the SNPs were of medium quality, with some ambiguity in differentiating between homozygotes and heterozygotes **(Figure 4B)**. Two other SNPs were rated as having inconclusive quality due to their sensitivity to DNA concentration and the resultant difficulty in scoring the marker genotypes **(Figure 4C),** while one SNP did not form scorable clusters and was regarded as having a bad quality (**Figure 4D**). A detailed report on the quality assessment of each SNP is presented in **Table 4**. SNPs that were easy to score formed three genotype clusters representing the two Mendelian homozygotes and one heterozygote. Those with inconclusive results formed either one or two genotype clusters, while the bad-quality SNPs did not create any meaningful clusters and were considered failed SNPs **(Table 4)**. Overall, the performance of 14 out of the 17 SNPs was deemed acceptable and selected as candidates for further validation.

**Figure 4.**
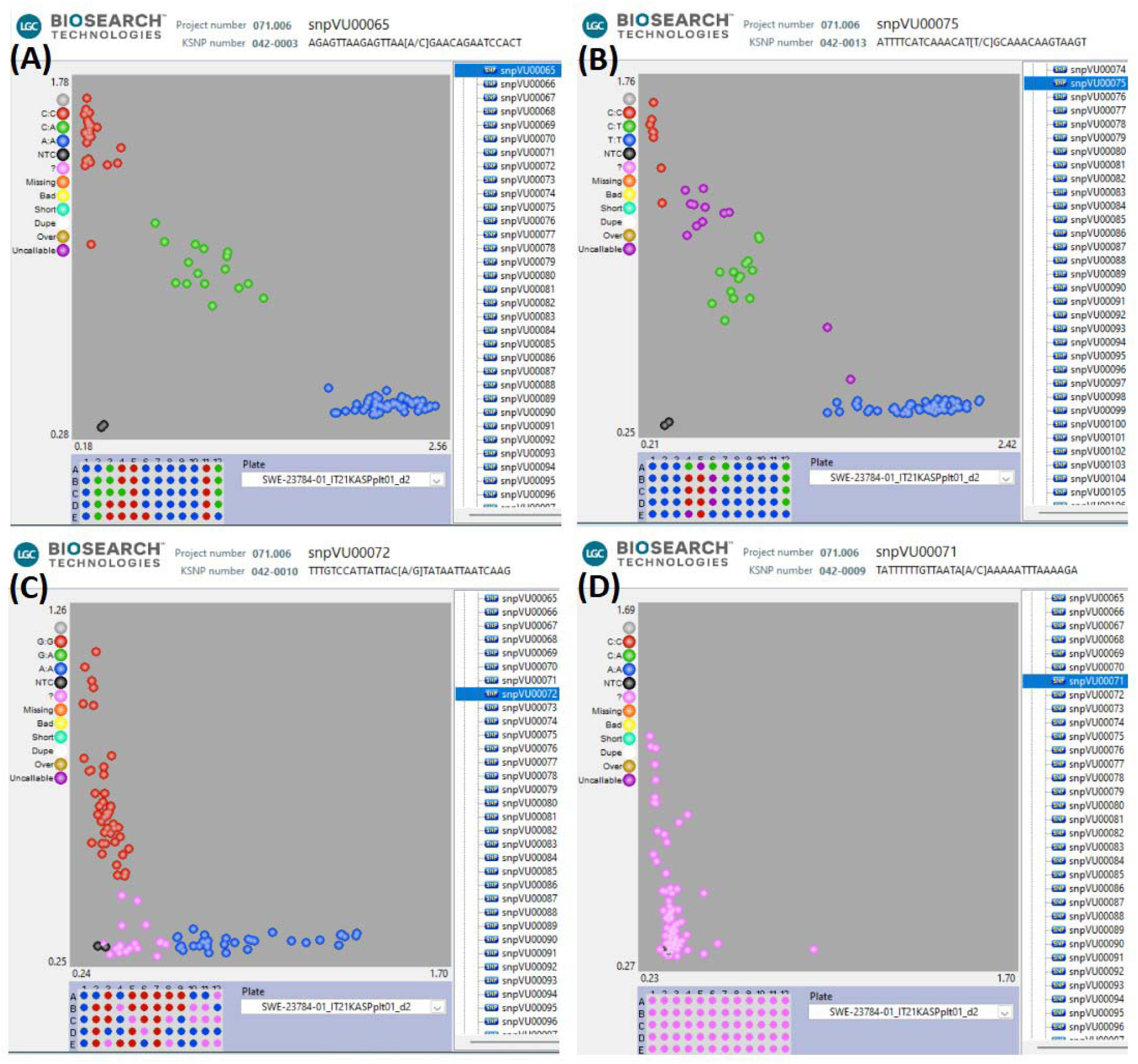
Visualizations of KASP assays technical verification of the candidate SNPs. **(A)** represents the 12 SNPs that had good quality KASP assays, **(B)** represents the 2 SNPs that had medium KASP assay quality and were still scorable, **(C)** represents the 2 SNPs that had inclusive results (only one or two genotype clusters), **(D)** the SNP that could not be scored since no clear genotype clusters were formed. In this figure, the blue and red clusters are the two Mendelian homozygous genotypes, while the green cluster is the heterozygous genotype. The black data points represent the no template controls (NTC). The pink data points marked with "?" represent samples that did not generate consistent signals or failed to amplify. The DNA sample plate layout is shown below the cluster plot.

**Table 4.** KASP assay technical verification report for 17 candidate SNPs for Striga resistance.

| CHR | SNP ID | KASP ID | SNP quality | Number of Clusters | Comment |
| --- | --- | --- | --- | --- | --- |
| Vu02 | 2_05789 | snpVU00063 | +++ | 3 | Easy to score |
| Vu02 | 2_18924 | snpVU00064 | +++ | 3 | Easy to score |
| Vu02 | 2_18925 | snpVU00065 | +++ | 3 | Easy to score |
| Vu02 | 2_22778 | snpVU00066 | +++ | 3 | Easy to score |
| Vu02 | 2_48732 | snpVU00067 | +++ | 3 | Easy to score |
| Vu07 | 2_00674 | snpVU00068 | +++ | 3 | Easy to score |
| Vu07 | 2_14573 | snpVU00069 | +++ | 3 | Easy to score |
| Vu07 | 2_20936 | snpVU00070 | +++ | 3 | Easy to score |
| Vu07 | 2_24995 | snpVU00071 | -- | 0 | No clusters |
| Vu07 | 2_27836 | snpVU00072 | - | 2 | Sensitive to DNA concentration |
| Vu10 | 2_32524 | snpVU00073 | - | 1 | Not scorable |
| Vu10 | 2_46726 | snpVU00074 | ++ | 3 | Lower DNA concentration migrates slower |
| Vu11 | 2_07557 | snpVU00075 | + | 3 | One extra HET subcluster, ambiguous scoring |
| Vu11 | 2_15481 | snpVU00076 | +++ | 3 | Easy to score |
| Vu11 | 2_42259 | snpVU00077 | ++ | 3 | HET cluster close to C:C cluster |
| Vu11 | 2_49024 | snpVU00078 | +++ | 3 | Easy to score |
| Vu11 | 2_50655 | snpVU00079 | + | 3 | HET cluster very close to T:T cluster |

+++ denotes very good quality SNP that is working well and is easy to score; ++ indicates good quality SNP that is working reasonably well but is sensitive to DNA concentration or clusters are close together, hence faulty annotations and less than 95% data recovery can occur; + refers to medium SNP quality that is working well but is difficult to score with confidence and/or many unamplified or un-callable data points, faulty annotation and less than 95% data recovery is expected; - denotes inconclusive results with only one or two clusters of data points identified (2x Hom and 1x Het is not present in the tested population); -- indicates bad SNP, this SNP is not working and no clusters are formed.

### 3.5 KASP marker validation

The validation exercise involved screening 20 cowpea genotypes in an artificially Striga-infested nursery and comparing the Striga phenotypic data with the marker genotypes on these cowpea genotypes. The phenotypic distribution of the 20 cowpea genotypes in response to Striga infestation is presented in **Figures 5A and B**. The resistant cowpea genotypes had zero Striga emergence, as portrayed by zero mean and median. In contrast, the susceptible cowpea genotypes showed dispersed distributions with the mean and median Striga count clearly above zero **(Figure 5A and B)**.

**Figure 5.**
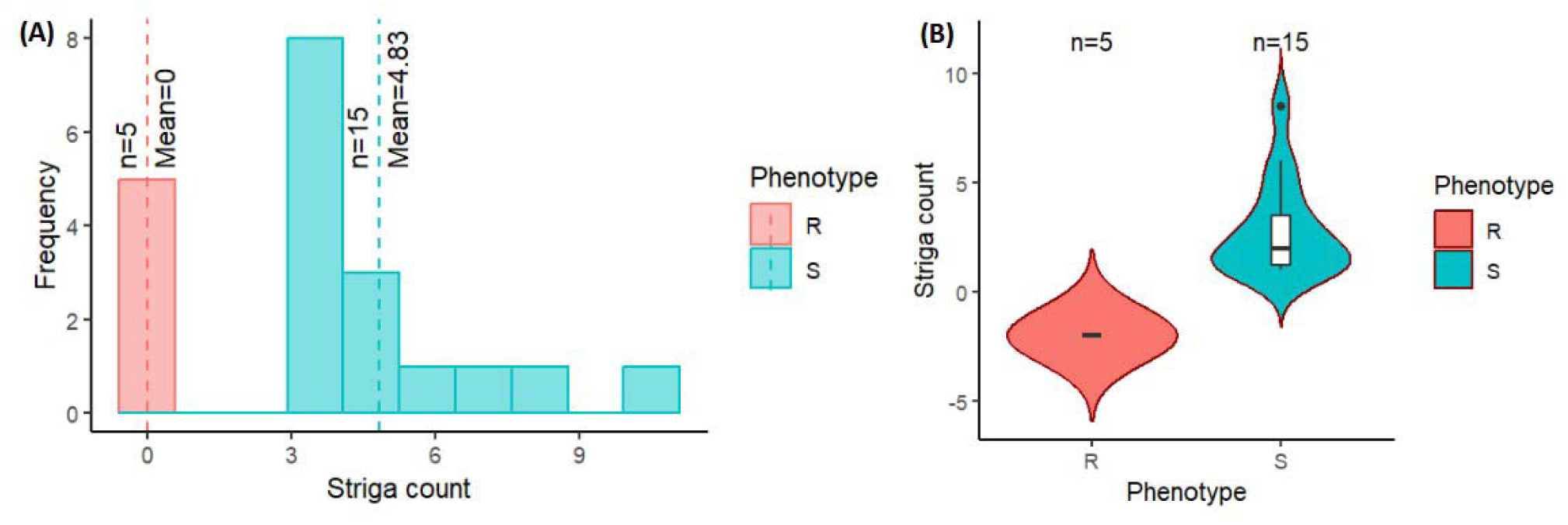
Distribution of Striga counts for the 20 cowpea genotypes evaluated in an artificially Striga-infested nursery at IITA Kano, Nigeria, grouped by their phenotypic status. **(A)** histogram showing the distribution of the resistant genotypes (R) in red and susceptible genotypes (S) in blue. The red and blue vertical dash lines depict the mean of the resistant and susceptible groups, respectively. **(B)** boxplot portraying the resistant genotypes (R) dispersion in red and susceptible genotypes (S) in blue. The black line at the center of the boxplot indicates the median Striga count.

Consequently, the counts of Striga emergence as influenced by the 20 cowpea genotypes were significantly different, with each genotype recording varying mean Striga count **(Table 5)**. Five cowpea genotypes were entirely immune to Striga (no Striga emerged in these plots), while the other genotypes differed in the average number of Striga ranging from 3-10.5, indicating different response levels to Striga **(Table 5)**. As expected, an old variety named Achishiru registered the highest number of Striga, followed by the landraces TVu-867 and Vita7, among the susceptible genotypes compared to low Striga counts registered in some of the released varieties like IT07K-318-33, IT98K-1111-1, IT08K-150-12, IT89KD-288 and IT98K-589-2 (Table 4), suggesting that these are Striga tolerant genotypes.

**Table 5.** Reactions of 20 selected cowpea genotypes to Striga infestation, evaluated in an artificially infested Striga nursery at IITA Kano station, Nigeria.

| Genotype | Rep I | Rep II | Mean Striga count | sd | Striga P/A Status |
| --- | --- | --- | --- | --- | --- |
| Achishiru | 15 | 6 | 10.50a | 6.36 | S |
| Tvu-8671 | 10 | 6 | 8.00ab | 2.83 | S |
| Vita7 | 9 | 5 | 7.00abc | 2.83 | S |
| Tvu-801 | 5 | 7 | 6.00bcd | 1.41 | S |
| IT86D-1010 | 5 | 5 | 5.00bcd | 0.00 | S |
| Tvu-1727 | 5 | 5 | 5.00bcd | 0.00 | S |
| Danila | 5 | 4 | 4.50bcd | 0.71 | S |
| TVX3236 | 4 | 4 | 4.00cd | 0.00 | S |
| CB27 | 6 | 1 | 3.50cde | 3.54 | S |
| IT07K-318-33 | 5 | 2 | 3.50cde | 2.12 | S |
| IT98K-1111-1 | 5 | 2 | 3.50cde | 2.12 | S |
| IT08K-150-12 | 3 | 3 | 3.00de | 0.00 | S |
| IT89KD-288 | 4 | 2 | 3.00de | 1.41 | S |
| IT98K-589-2 | 4 | 2 | 3.00de | 1.41 | S |
| Sanzi | 5 | 1 | 3.00de | 2.83 | S |
| B301 | 0 | 0 | 0.00e | 0.00 | R |
| IT13K-1308-5 | 0 | 0 | 0.00e | 0.00 | R |
| IT97K-499-35 | 0 | 0 | 0.00e | 0.00 | R |
| IT99K-573-1-1 | 0 | 0 | 0.00e | 0.00 | R |
| IT99K-573-2-1 | 0 | 0 | 0.00e | 0.00 | R |
| Error mean square |  |  | 3.26 |  |  |
| Error degrees of freedom (DF) |  |  | 19.00 |  |  |
| LSD |  |  | 3.78 |  |  |

Rep I and Rep II represent Striga counts from replications one and two, respectively; sd is the standard deviation; S and R represent each line's susceptible and resistant status, respectively based on striga presence-absence (P/A) assessment. The means of genotypes with the same letter are not significantly different.

Using a chi-square test of independence, the study compared the association between each of the 14 candidate SNPs with the phenotypic status of the 20 cowpea genotypes (**Table 6).** Five SNPs in close proximity to the major association signal on chromosome Vu11 displayed highly significant deviations (P ≤ 0.01) from the null expectation of no marker–phenotype correlation, supporting their strong association with genomic regions influencing Striga resistance in cowpea **(Table 6)**. Five other SNPs positioned close to each other on chromosome Vu02 showed moderate statistical significance (P ≤ 0.05). The rest of the SNPs on Chromosomes Vu07 and Vu10 revealed no significant chi-square values **(Table 6).**

**Table 6.** Chi-square test of independence between the candidate SNPs and the Striga resistance in cowpea.

| CHR | SNP ID | KASP ID | Allele | Obs(R,S) | Exp(R,S) | N | DF | $\chi^2$ | p-value | Decision ($\alpha \leq 0.05$) |
| --- | --- | --- | --- | --- | --- | --- | --- | --- | --- | --- |
| Vu02 | 2_05789 | snpVU00063 | AA | 1,11 | 3.00,9.00 | 20 | 1 | 4.44 | 0.035 | * |
|  |  |  | GG | 4,4 | 2.00,6.00 |  |  |  |  |  |
| Vu02 | 2_18924 | snpVU00064 | CC | 4,4 | 2.00,6.00 | 20 | 1 | 4.44 | 0.035 | * |
|  |  |  | TT | 1,11 | 3.00,9.00 |  |  |  |  |  |
| Vu02 | 2_18925 | snpVU00065 | AA | 1,11 | 3.00,9.00 | 20 | 1 | 4.44 | 0.035 | * |
|  |  |  | CC | 4,4 | 2.00,6.00 |  |  |  |  |  |
| Vu02 | 2_22778 | snpVU00066 | AA | 4,4 | 2.00,6.00 | 20 | 1 | 4.44 | 0.035 | * |
|  |  |  | GG | 1,11 | 3.00,9.00 |  |  |  |  |  |
| Vu02 | 2_48732 | snpVU00067 | GG | 1,11 | 3.00,9.00 | 20 | 1 | 4.44 | 0.035 | * |
|  |  |  | TT | 4,4 | 2.00,6.00 |  |  |  |  |  |
| Vu07 | 2_00674 | snpVU00068 | AA | 5,12 | 4.25,12.75 | 20 | 1 | 1.18 | 0.278 | Ns |
|  |  |  | GG | 0,3 | 0.75,2.25 |  |  |  |  |  |
| Vu07 | 2_14573 | snpVU00069 | AA | 5,12 | 4.25,12.75 | 20 | 1 | 1.18 | 0.278 | Ns |
|  |  |  | GG | 0,3 | 0.75,2.25 |  |  |  |  |  |
| Vu07 | 2_20936 | snpVU00070 | GG | 5,12 | 4.25,12.75 | 20 | 1 | 1.18 | 0.278 | Ns |
|  |  |  | TT | 0,3 | 0.75,2.25 |  |  |  |  |  |
| Vu10 | 2_46726 | snpVU00074 | AA | 1,10 | 2.75,8.25 | 20 | 1 | 1.18 | 0.069 | Ns |
|  |  |  | GG | 4,5 | 2.25,6.75 |  |  |  |  |  |
| Vu11 | 2_07557 | snpVU00075 | CC | 4,1 | 1.25,3.75 | 20 | 1 | 10.76 | 0.001 | *** |
|  |  |  | TT | 1,14 | 3.75,11.25 |  |  |  |  |  |
| Vu11 | 2_15481 | snpVU00076 | GG | 4,1 | 1.25,3.75 | 20 | 1 | 10.76 | 0.001 | *** |
|  |  |  | TT | 1,14 | 3.75,11.25 |  |  |  |  |  |
| Vu11 | 2_42259 | snpVU00077 | CC | 1,15 | 4.00,12.00 | 20 | 1 | 15 | 0.0001 | *** |
|  |  |  | GG | 4,0 | 1.00,3.00 |  |  |  |  |  |
| Vu11 | 2_49024 | snpVU00078 | AA | 4,2 | 1.50,4.50 | 20 | 1 | 7.94 | 0.0048 | ** |
|  |  |  | GG | 1,13 | 3.50,10.50 |  |  |  |  |  |
| Vu11 | 2_50655 | snpVU00079 | CC | 5,3 | 2.00,6.00 | 20 | 1 | 10 | 0.0016 | ** |
|  |  |  | TT | 0,12 | 3.00,9.00 |  |  |  |  |  |

CHR refers to chromosome; SNP ID is the original name of the SNP marker; KASP ID is the new name of the candidate SNPs after developing the KASP assay; Obs(R, S) refers to the observed number of resistant and susceptible cowpea genotypes; Exp(R, S) refers to the expected number of resistant and susceptible cowpea genotypes; N is the total number of cowpea genotypes tested; DF is the degrees of freedom, $\chi^2$ is the chi-square statistic, the p-value is the probability value associated with the chi-square test.

Further efforts to group the phenotypic reactions by marker alleles revealed a strong marker-phenotype association for the candidate markers on chromosome Vu11 and a weak relation for the other SNPs **(Figure 6, Supplementary Figure 1A,B,C,D)**. For instance, the GG allele of the marker snpVU00077 on Vu11 was responsible for 100% of the cowpea genotypes that had a Striga-resistant phenotype, while the alternative allele CC was carried by 94% of phenotypically susceptible genotypes, with only 6% phenotypic misclassification **(Figure 6).** This observation was supported by a highly significant chi-square test and high Cramer’s correlation (V _Cramer_ =0.86). Similar results were observed for all five proximal SNPs on Vu11 **(Supplementary Figure 1D),** suggesting that these markers are strongly associated with genes underlying Striga resistance in cowpea. However, a relatively high percentage of misclassifications and V _Cramer_ less than 0.5 were observed for the remaining candidate markers on chromosomes Vu02, Vu07, and Vu10 **(Figure 6, Supplementary Figure 1A,B,C),** suggesting weak marker-trait association.

**Figure 6.**
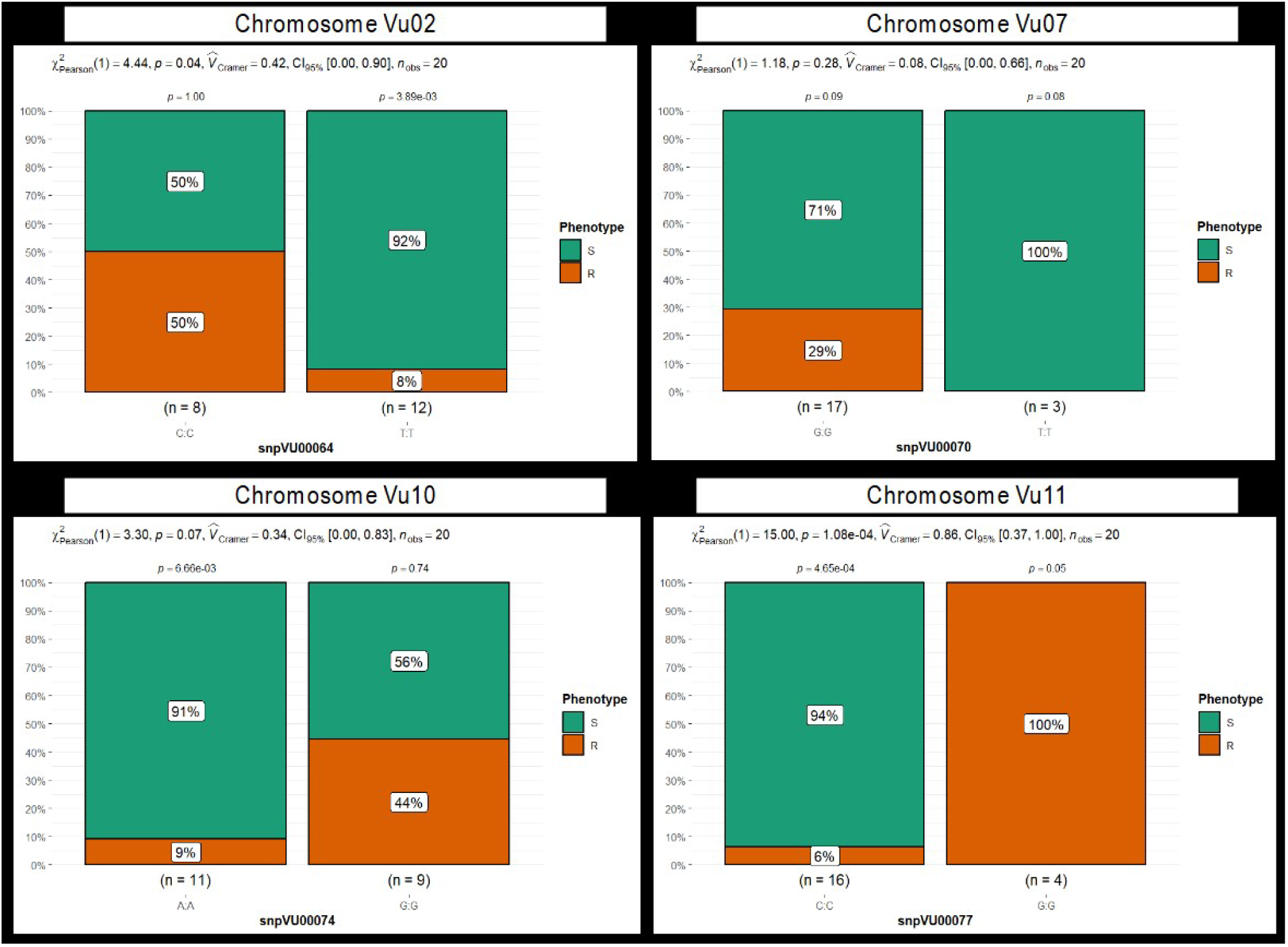
Stacked bar chart depicting the grouping of phenotypic reactions to Striga by marker alleles. Four markers: snpVU00064, snpVU00070, snpVU00074, and snpVU0007, represent the candidate regions on chromosomes Vu02, Vu07, Vu10, and Vu11, respectively. Results for all 14 markers are presented in **Supplementary figure 1**. The 20 cowpea genotypes are grouped based on their phenotypic reaction to Striga into resistant (R) and susceptible (S) classes, and the stacked bar charts are plotted to reflect the percentage of genotypes in each phenotypic category that carry the two alleles of each SNP marker. Summary statistics are presented at the top of each stacked bar chart that includes the Pearson chi-square test (χ^2^_Pearson_), probability (p) presented for each marker allele class, and for the overall chi-square test of independence, effect size measured by Cramer’s correlation (V □_Cramer_), 5% confidence interval (CI_95%_) and number of observations (n_obs_).

Similarly, a t-test was conducted to compare the differences in the mean Striga count between cowpea genotypes carrying the two alleles of each marker. Results for four markers representing the candidates on chromosomes Vu02, Vu07, Vu10, and Vu11 are presented in **Figure 7**. Highly significant differences (p ≤ 0.001) between the mean of the allelic groups were found only for the markers on chromosome Vu11, while the other markers were not significant, strongly supporting the chi-square test results. The marker alleles conferring resistance to Striga had significantly lower mean Striga counts than the alternative alleles. The results of the t-test for all 14 markers specifying the alleles underlying resistance and susceptibility and how they differ have been presented in **supplementary Figure 2A, B, C, D.**

**Figure 7.**
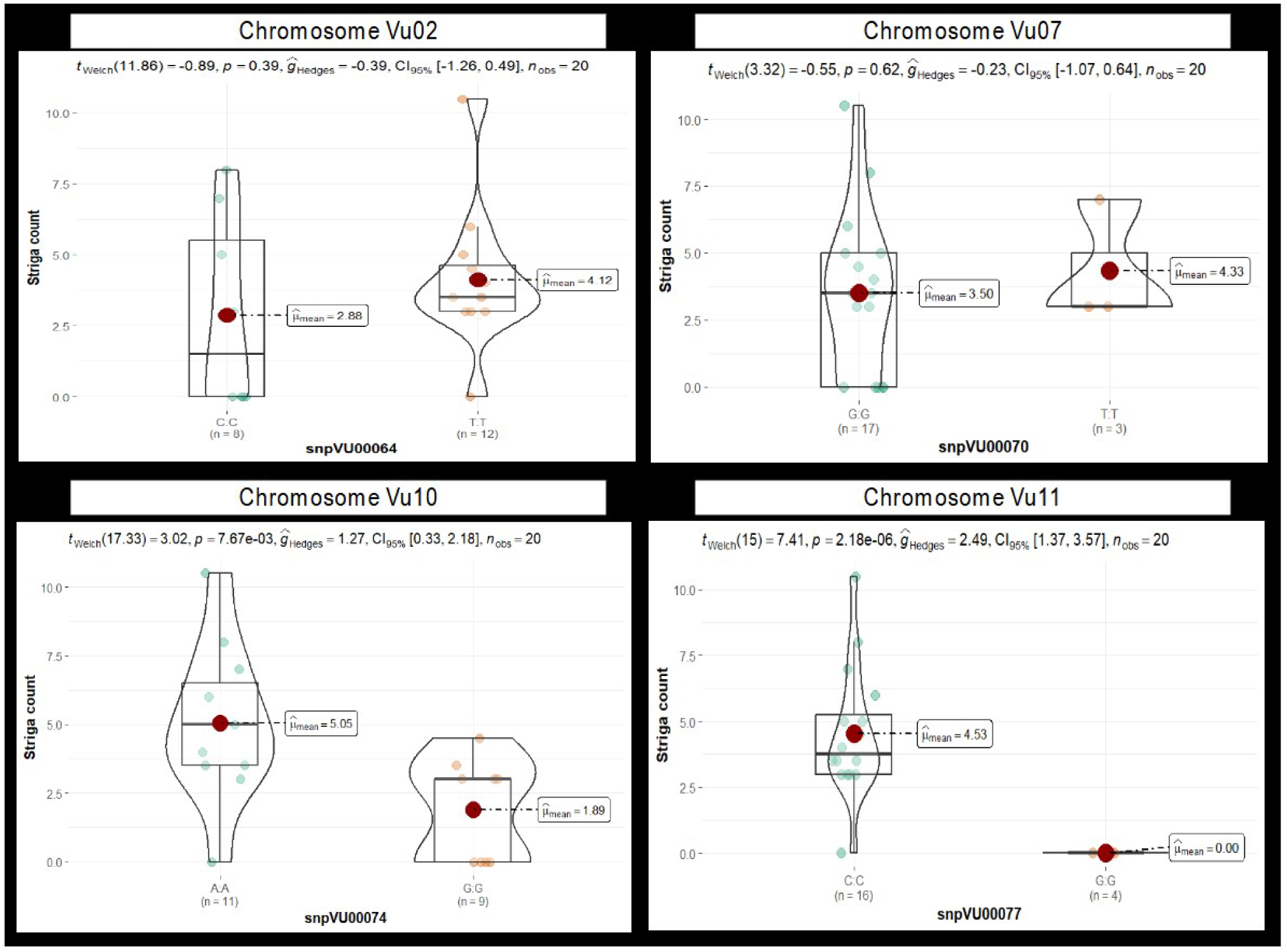
The violin chart depicting the difference in mean Striga counts between cowpea genotypes carrying the two alleles of each candidate marker. Results of four markers, snpVU00064, snpVU00070, snpVU00074, and snpVU0007, are presented to represent the candidate regions on chromosomes Vu02, Vu07, Vu10, and Vu11, respectively. Results for all 14 markers are presented in **Supplementary figure 2**. The mean (µ ^_mean_) of each allelic group is indicated by a red dot. Summary statistics are presented at the top of the violin chart, including t-statistical test (t_welch_), probability (p) for the t-test, effect size measured by Hedges’ g (ĝ_Hedges_), a 5% confidence interval (CI_95%_) and number of observations (n_obs_).

## 4 Discussion

*Striga gesnerioides*, commonly known as cowpea witchweed, is a parasitic plant that poses a significant threat to cowpea (*Vigna unguiculata*) production, particularly in West and Central Africa. This parasitic weed attaches to cowpea roots, extracting essential nutrients and water, leading to substantial yield reductions and, in severe infestations, complete crop failure (Li et al., 2009; Sadda et al., 2021). The multiplication of *S. gesnerioides* is facilitated by its prolific seed production, with a single capsule containing hundreds of seeds that can remain viable in the soil for several years (Mohamed et al., 2001). Addressing the challenges posed by *S. gesnerioides* requires integrated management strategies, among which developing and cultivating Striga-resistant cowpea varieties is pivotal (Abdullahi et al., 2022). The application of molecular markers has been emphasized as the best approach to facilitate breeding for resistance to this parasitic weed, given its race complexity (Ouédraogo et al., 2001; Boukar et al., 2004; Li et al., 2009; Ouédraogo et al., 2012; Essem et al., 2019). Successfully deploying markers in breeding requires discovering and validating markers that closely tag the resistance loci. In the case of *S. gesnerioides*, which has multiple races (Botanga and Timko, 2006; Ohlson and Timko, 2020), it is vital to develop molecular markers linked to resistance against each race. These markers would facilitate the pyramiding of resistance genes, hence the development of cowpea varieties suitable for cultivation across the Striga-affected area of WA. The present study significantly contributes to these efforts by uncovering key loci and developing molecular markers underlying resistance to Nigeria’s dominant Striga race SG3.

### 4.1 Genomic regions associated with Striga resistance

The present study deciphered genomic regions controlling *S. gesnerioides* resistance in cowpea by utilizing a high-density single nucleotide polymorphism marker and diverse cowpea minicore genotypes. The phenotypic data from the two test sites revealed significant genetic variation in resistance to *Striga gesnerioides* with skewed distribution, indicating that major-effect genes likely govern resistance. The finding aligns with prior studies, which suggested that resistance to *S. gesnerioides* in cowpea is often controlled by specific resistance (R) genes exhibiting major effects. Notably, the cowpea genotypes B301 and IT82D-849 exhibited resistance controlled by a single dominant gene effective against Striga races SG1, SG2, and SG3 found in Nigeria (Singh and Emechebe, 1990; Atokple et al., 1993; Li et al., 2009). In a cross of cowpea genotypes HTR (from Niger) and Wango-1 (from Burkina Faso), it was reported that resistance to *S. gesnerioides* race SG1 in HTR was controlled by one or two dominant genes that are non-allelic to the genes in B301 and IT82D-849 (Timko et al., 2007; Li et al., 2009). In addition, monogenic but recessive inheritance of resistance to *S. gesnerioides* race SG3 from Niger was also reported (Touré et al., 1997). However, the interaction between cowpea and *S. gesnerioides* is complex due to the existence of multiple parasite races. Seven distinct races (SG1 to SG7) were identified, each capable of overcoming specific resistance genes in cowpea (Li et al., 2009; Ohlson and Timko, 2020). This variability necessitates discovering key loci involved, followed by stacking multiple resistance genes to achieve broad-spectrum and durable resistance.

A genome-wide association analysis revealed a major association signal on chromosome Vu11, alongside f significant minor signals on chromosomes Vu02, Vu07, and Vu10 linked to Striga resistance. The prominent Vu11 signal aligns closely with the a coiled-coil nucleotide-binding site leucine-rich repeat (CC-NBS-LRR) protein, encoded by the RSG3-301 gene, was implicated in race-specific resistance to *S. gesnerioides* in cowpea (Li and Timko, 2009). This proximity suggests that similar R-genes may underlie the Vu11 association signal.. Additionally, a BLAST analysis using primers from earlier marker technologies (e.g., AFLP, SSR, and SCAR) returned best matches within the same Vu11 region **(Table 7)**, further supporting the colocalization of historical markers with our GWAS-identified signals.. The previously reported genes spanning the same area on chromosome Vu11 included *Rsg3* (conferring resistance to Striga race *SG3* and SG5)*, Rsg2–1* (effective against Striga race SG1), and *Rsg4–3* (effective against Striga race SG3) (Ouédraogo et al., 2001; Essem et al., 2019). Earlier linkage mapping studies identified resistance loci to Striga in cowpea via bi- parental populations. Ouédraogo et al. (2002) mapped a race-specific resistance gene (designated *Rsg3* and *Rsg994*) to linkage group 1 (equivalent to chromosome Vu10 in the reference genome), conferring resistance to Striga race SG1. Boukar et al. (2004) mapped another resistance locus, Rsg1, to linkage group 1 (chromosome Vu08), conferring race-specific resistance to SG3 The studies above used traditional linkage mapping in F_2_ bi-parental populations to identify the Striga resistance loci. In comparing these classical mapping results with our GWAS findings, we observed a strong correspondence between previously reported loci and our newly identified association signals **(see Table 7)**. This concordance supports the involvement of these genomic regions, now validated by independent GWAS in mediating Striga resistance.

**Table 7.** Previously mapped Striga resistance QTL in cowpea with some of the loci occupying the same genomic region discovered in the present GWAS.

| Locus name | Striga race | Marker type | Marker name | LG | Dist(cM) | Population | BLAST search | References |
| --- | --- | --- | --- | --- | --- | --- | --- | --- |
| Rsg3 | SG3 | SSR | SSR-1 | - | - | IT97K-499-35 x SARC-LO2 (F8 RIL) | Vu11 (1,928,028 bp) | (Essem et al., 2019) |
| Rsg3 | SG3, SG5 | SCAR | C42-2B | - | - | IT97K-499-35 x SARC-LO2 (F8 RIL) | Vu11 (1,554,567 bp) | (Essem et al., 2019) |
| Rsg2-1 | SG1 | AFLP-SCAR | E-ACT/M-CAA<sub>524</sub> (**61RM2**) | 1 | 0.9 | Tvx 3236 x IT82D-849 (F2 population) | Vu11 (1,752,097 bp) | (Ouédraogo et al., 2001) |
| Rsg4-3 | SG3 | AFLP | E-ACA/M-CAT150 and E-AGC/M-CAT80 | 1 | 2.7-4.1 | Tvu 14676×IT84 S-2246-4 (F2 population) | Vu11<sup>#</sup> | (Ouédraogo et al., 2001) |
| Rsg3 | SG1 | AFLP | E-AGA/M-CTA460 and E-AGA/M-CAG300 | 6 | 2.5-2.6 | Tvx 3236 x Gorom-Suvita 2 (F2 population) | Vu10<sup>#</sup> | (Ouédraogo et al., 2002) |
| 994-Rsg | SG1 | AFLP | E-AAG/M-AAC450 and E-AAG/M-AAC150 | 6 | 2.0-2.1 | Tvx 3236 x IT81D-994 (F2 population) | Vu10<sup>#</sup> | (Ouédraogo et al., 2002) |
| Rsg1 | SG3 | AFLP/S CAR | E-ACT/M-CTC115 and E-ACT/M-CAC115 (**SEACTMC AC83/85**) | 1 | 3.2-4.8 | IT93K-693-2 x IAR1696 (F2 population) | Vu08 (25,734,040 bp) | (Boukar et al., 2004) |

<sup>#</sup>Blast search has not been conducted since the related AFLP markers are not cloned, however, LG1 and LG6 of Ouédraogo et al., (2002) has been matched to chromosomes Vu11 and Vu10 respectively of the cowpea reference genome in Phytozome (Muchero et al., 2009; Essem et al., 2019).

### 4.2 Candidate genes for Striga resistance

Through our candidate gene analysis, we identified 20 unique proteins that have been implicated in plant defense and immune response signaling, collectively annotated to 64 cowpea genes spanning the GWAS-identified association intervals.. Notably, among these is a leucine-rich repeat (LRR)- containing protein mapped near the major association peak on chromosome Vu11.LRR domains are well-established as pivotal in pathogen recognition and activation of plant defense responses, as seen in both extracellular receptor kinases and intracellular NB-LRR immune receptors (Jones and Jones, 1997; McHale et al., 2006). An earlier study in cowpea pinned leucine-rich repeat (CC-NBS-LRR) protein to a gene-for-gene resistance mechanism in the interactions between Striga and cowpea, with a corresponding gene in cowpea named *RSG3-301* (Li and Timko, 2009). The authors also unlocked the hypothesis of race specificity resistance by silencing the *RSG3-301* gene in cultivar B301, which rendered it susceptible to race RG3 but remained resistant to races SG2 and SG5. Another gene PTHR22952:SF183 - TRANSCRIPTION FACTOR TGA5 on Vu11, has been found to work along with TGA2 and TGA6, playing a crucial role in plant systemic acquired resistance (SAR), a broad-spectrum defense mechanism induced after a local infection by avirulent pathogens (Zhang et al., 2003). On chromosome Vu10, the likely genes involved in plant defense were ETHYLENE-RESPONSIVE TRANSCRIPTION FACTOR ERF003, WRKY DNA-binding domain, and PF03763 - Remorin, C-terminal region (Remorin_C). Ethylene-responsive transcription factors (ERFs) are integral components of the APETALA2/ERF superfamily, playing pivotal roles in regulating plant responses to biotic and abiotic stresses (Müller and Munné-Bosch, 2015). ERFs regulate molecular response to pathogen attack by binding to specific cis-acting elements in the promoters of stress-responsive genes, thereby modulating their expression (Müller and Munné-Bosch, 2015). WRKY transcription factors, on the other hand, have been associated with responses to various pathogens in cowpea, particularly exhibiting differential expressions when the plants were challenged with *Fusarium oxysporum*, a pathogen responsible for Fusarium wilt (Hao et al., 2024). This finding underscores the significant role of WRKY genes in cowpea’s defense mechanisms against biotic stresses. Given the conserved nature of WRKY transcription factors across plant species, it is plausible that manipulating specific WRKY genes in cowpea could enhance resistance to Striga. However, targeted studies are required to identify which WRKY genes are involved and to elucidate their mechanisms of action in the context of Striga resistance. In the context of plant defense, remorins have been observed to interact with receptor-like kinases (RLKs) and pathogen effectors, suggesting a role in the early stages of immune signaling (Yu, 2020). Chromosome Vu07 harbored genes encoding MADS-box transcription factors, cysteine-rich receptor-like kinases, MYB-like proteins, and GATA-type zinc finger transcription factors, all of which play roles in plant development and stress responses (Abdullah-Zawawi et al., 2021; Zhang et al., 2023a, 2023b). The genes on chromosome Vu02, included, among others, pentatricopeptide repeat (PPR) proteins, C2H2-like zinc finger proteins, glucuronosyltransferase PGSIP8-like, receptor-like kinases, and lipid transfer proteins that contribute to the complex network of plant defense mechanisms, each playing distinct roles in responding to biotic and abiotic stresses (Goff and Ramonell, 2007; Finkina et al., 2016; Zhang et al., 2020; Huang et al., 2022; Meng et al., 2024).

### 4.3 Marker development

Seventeen markers tagging GWAS-identified association intervals for Striga resistance were converted into KASP assays and independently validated across 20 cowpea genotypes with known resistance profiles. Five of these markers, all situated near the major association signal on chromosome Vu11, were confirmed to be associated with resistance.. KASP is a globally recognized technology for SNP genotyping, known for being user-friendly, relatively cheap, and can be automated, ensuring high throughput genotyping (Ongom et al., 2021; Dipta et al., 2024). The five Striga-associated SNP markers designated by Intertek KASP assay IDs; snpVU00075, snpVU00076, snpVU00077, snpVU00078, and snpVU00079 are all positioned proximally to each other within the major QTL region on Vu11. Previously reported marker systems for Striga resistance were mostly AFLPs, SCARs, and SSRs (Ouédraogo et al., 2001, 2002; Boukar et al., 2004; Essem et al., 2019). Although some of these old marker technologies were reportedly effective in marker-assisted selection (MAS) for Striga resistance (Larweh et al., 2017; Omoigui et al., 2017), they have limitations, including labor-intensive gel electrophoresis or capillary electrophoresis, lack of automation and cumbersomeness for high-throughput applications, high cost per data point, and low reproducibility and accuracy (Mut et al., 2008; Dipta et al., 2024). The present study underscores the potential of new KASP-based SNP markers for efficient MAS in cowpea breeding programs aimed at developing *Striga*-resistant varieties.

## 5 Conclusion

This study deepens our understanding of the genetic architecture of *Striga* resistance in cowpea by pinpointing key association signals and candidate genes.. A dominant signal on chromosome Vu11 aligns with a known leucine-rich repeat (LRR) resistance gene, reinforcing our confidence in targeting this region for breeding applications.. SNPs within these associated intervals were converted to breeder-friendly KASP markers for routine breeding applications. These markers, particularly those on chromosome Vu11, provide valuable tools for breeding programs focused on developing *Striga*-resistant cowpea varieties, thereby contributing to improved cowpea productivity in *Striga*-infested regions. Ongoing efforts aim to validate these markers across diverse genetic backgrounds using segregating bi-parental populations to broaden their applicability. Since multiple Striga races affect cowpea across West Africa, pyramiding resistance alleles will be essential for achieving durable, broad-spectrum resistance. Continued research is also needed to identify and validate resistance loci against other predominant Striga races.

## 7 Author Contributions

Conceptualization, P.O.O.; methodology, P.O.O.; software, P.O.O.; validation, P.O.O., O.B, and C.F.; formal analysis, P.O.O; investigation, P.O.O. O.B, and C.F; resources, O.B. and C.F.; data curation, P.O.O.; writing—original draft preparation, P.O.O.; writing—review and editing, P.O.O., O.B and C.F.; visualization, P.O.O.; supervision, O.B, and C.F.; project administration, O.B.; funding acquisition, O.B. All authors have read and agreed to the published version of the manuscript.

## 9 Acknowledgments

We wish to express our sincere gratitude to the International Institute of Tropical Agriculture (IITA) for graciously hosting this study and granting access to their phenotyping facilities. This support was instrumental in enabling the thorough evaluation of cowpea-*Striga gesnerioides* interactions using the minicore collection, which IITA maintains as part of its global germplasm repository. We also wish to thank the team at the University of California, Riverside (UCR) for genotyping the cowpea minicore population using the high-density iSelect SNP array and making this valuable data publicly available at https://doi.org/10.1002/leg3.95. This open-access resource, covering over 51,000 SNP loci, greatly facilitated our genome-wide association analyses. Finally, we extend our appreciation to all field technicians and support personnel involved in data generation and analysis. Their collective commitment has been central to the discovery and development of KASP markers, which will underpin future marker-assisted breeding efforts for Striga-resistant cowpea varieties.

## 12 Data Availability Statement

SNP data was previously published: María Muñoz-Amatriaín et al 2021. Legume Science. https://doi.org/10.1002/leg3.95. All other data reported in this study has been provided as supplementary materials.

