## Supplementary material for "Discovery and development of single-nucleotide polymorphism markers for resistance to *Striga gesnerioides* in cowpea (*Vigna unguiculata*)": Supplementary_Figures.docx

### 1.1 Supplementary Figure 1

Chromosome Vu02

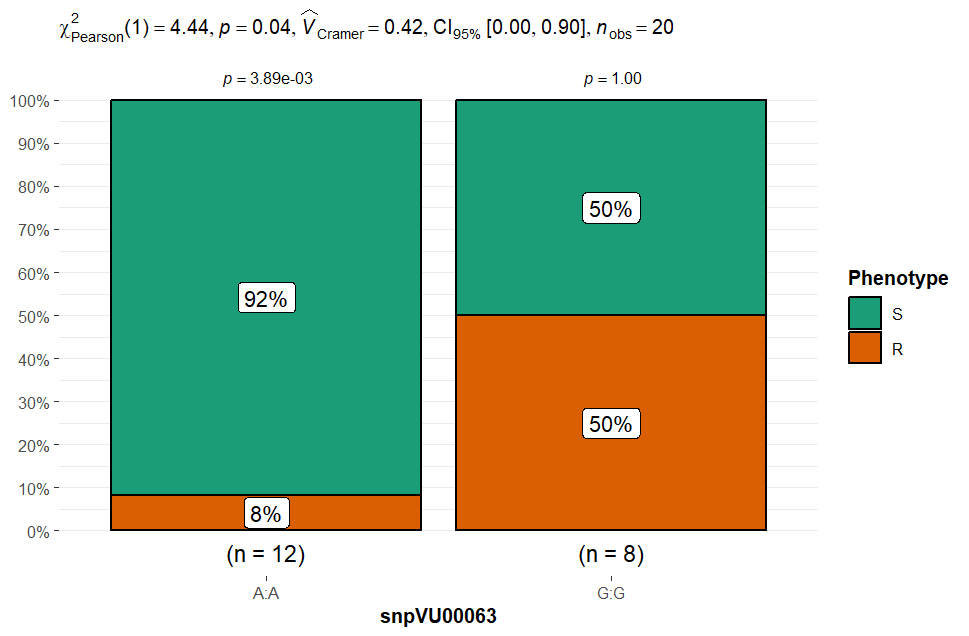

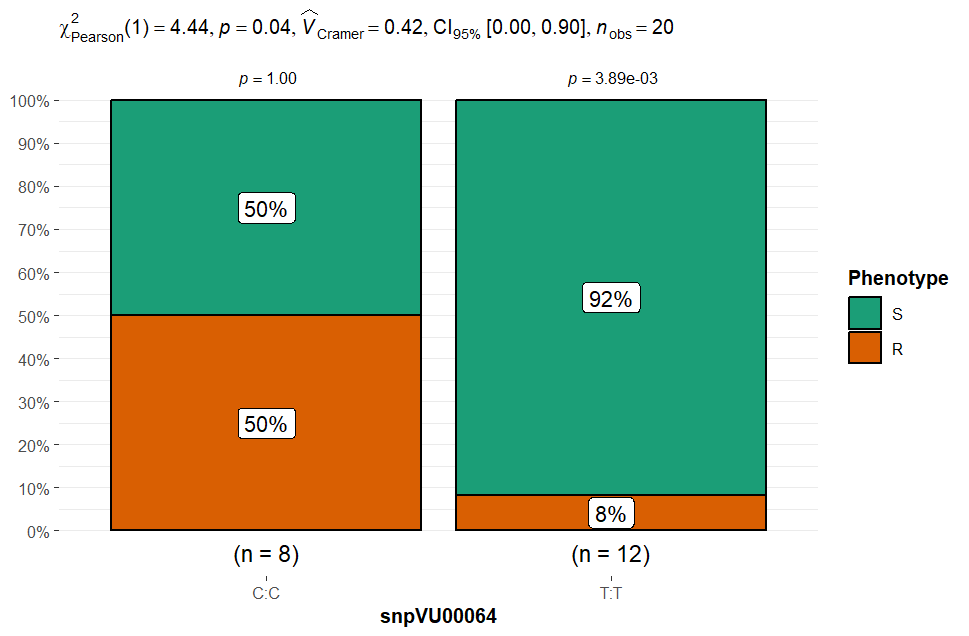

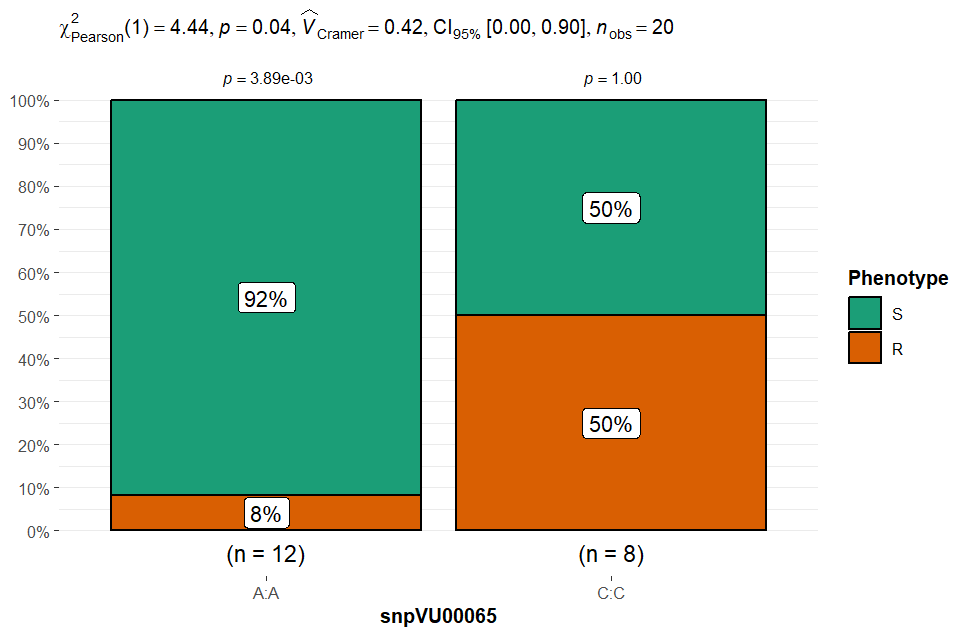

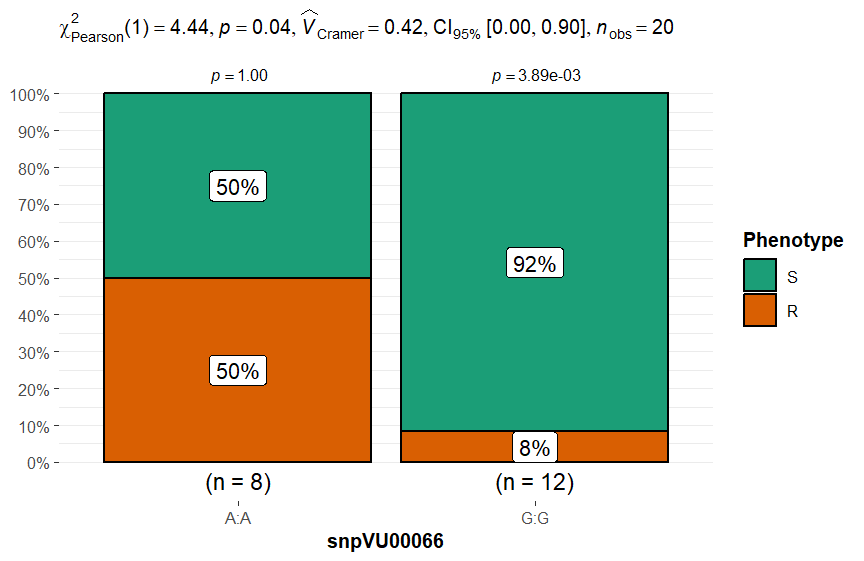

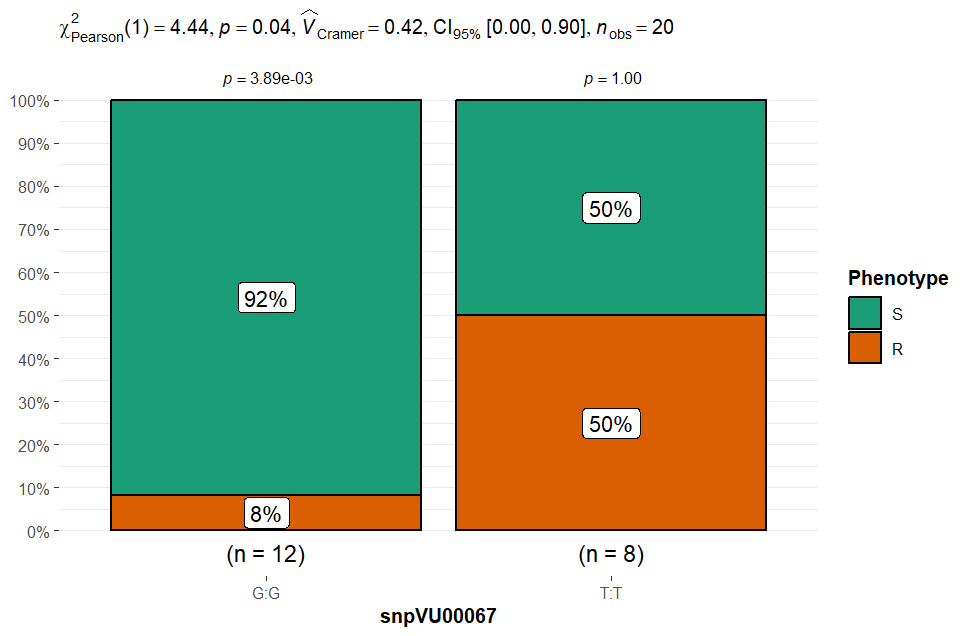

**Supplementary Figure 1A.** Chi-square test results for marker validation presented as stacked bar charts depicting the grouping of phenotypic reactions to Striga by the alleles five proximal candidate markers on chromosome Vu02. The 20 cowpea lines are grouped based on their phenotypic reaction to striga into resistant (R) and susceptible (S) classes and the stacked bar charts are plotted to reflect the percentage of lines in each phenotypic category that carry the two alleles of a SNP marker. Summary statistics are presented at the top of each stacked bar chart that include chi square test (χ^2^), probability (p) presented for each marker allele class and for overall test of chi square test of independence, Cramer’s correlation (V̂_Cramer_), 5% confidence interval (CI_95%_) and number of observations (n_obs_).

Chromosome Vu07 and Vu10

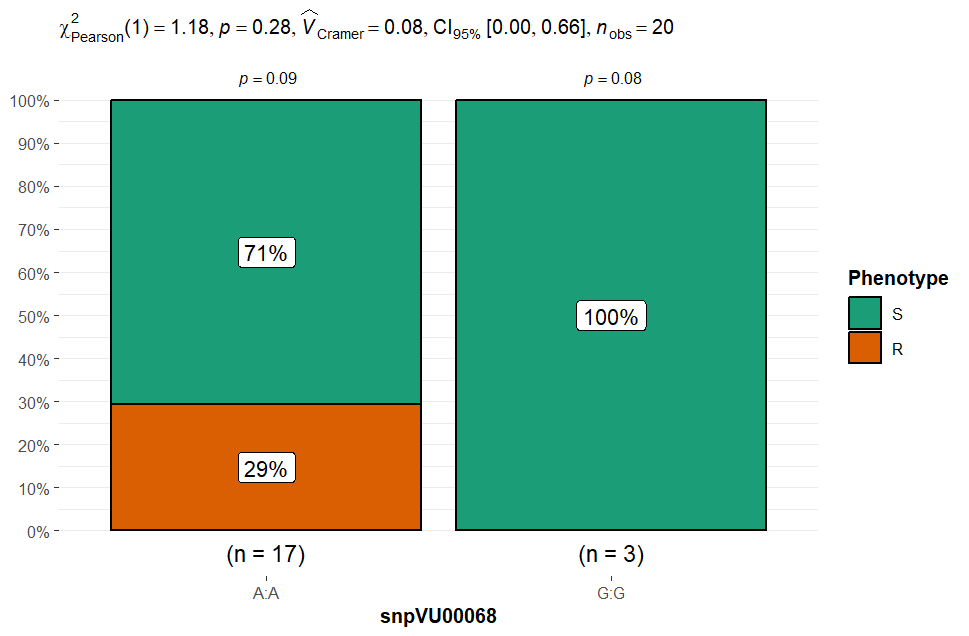

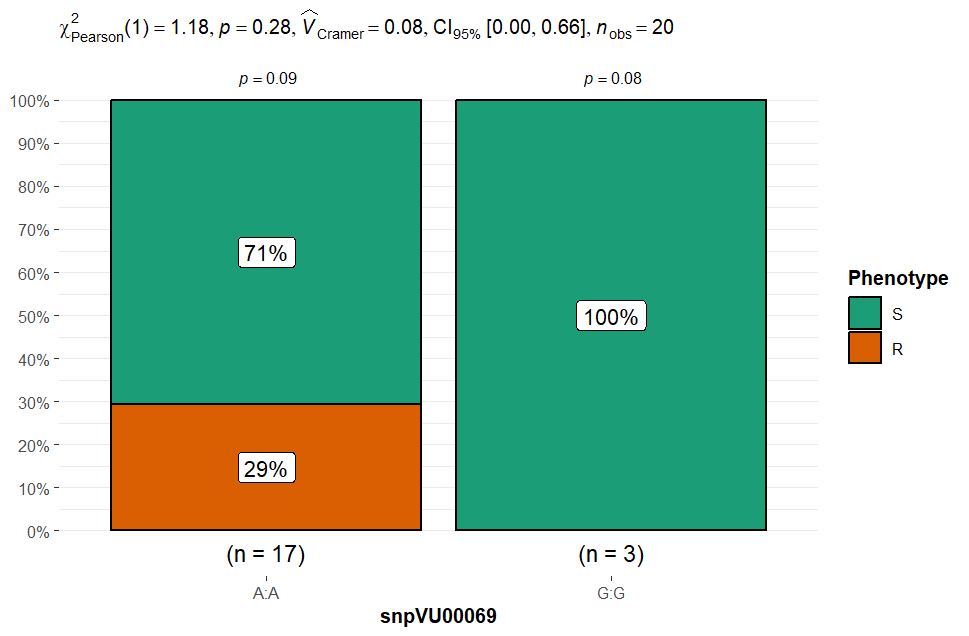

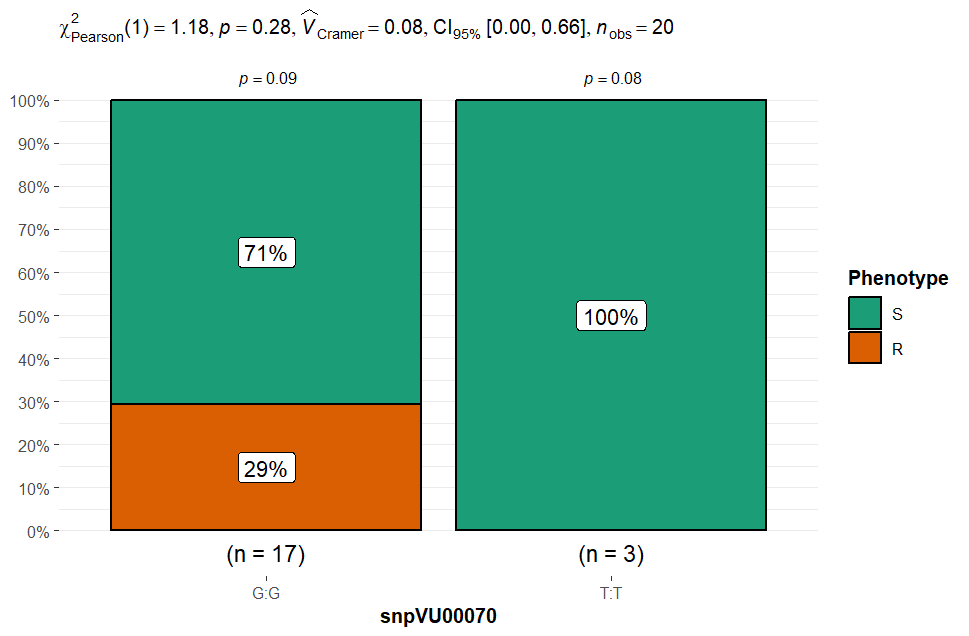

**Supplementary Figure 1B.** Chi-square test results for marker validation presented as stacked bar charts depicting the grouping of phenotypic reactions to Striga by the alleles three proximal candidate markers on chromosome Vu07. The 20 cowpea lines are grouped based on their phenotypic reaction to striga into resistant (R) and susceptible (S) classes and the stacked bar charts are plotted to reflect the percentage of lines in each phenotypic category that carry the two alleles of a SNP marker. Summary statistics are presented at the top of each stacked bar chart that include chi square test (χ^2^), probability (p) presented for each marker allele class and for overall test of chi square test of independence, Cramer’s correlation (V̂_Cramer_), 5% confidence interval (CI_95%_) and number of observations (n_obs_).

Chromosome Vu10

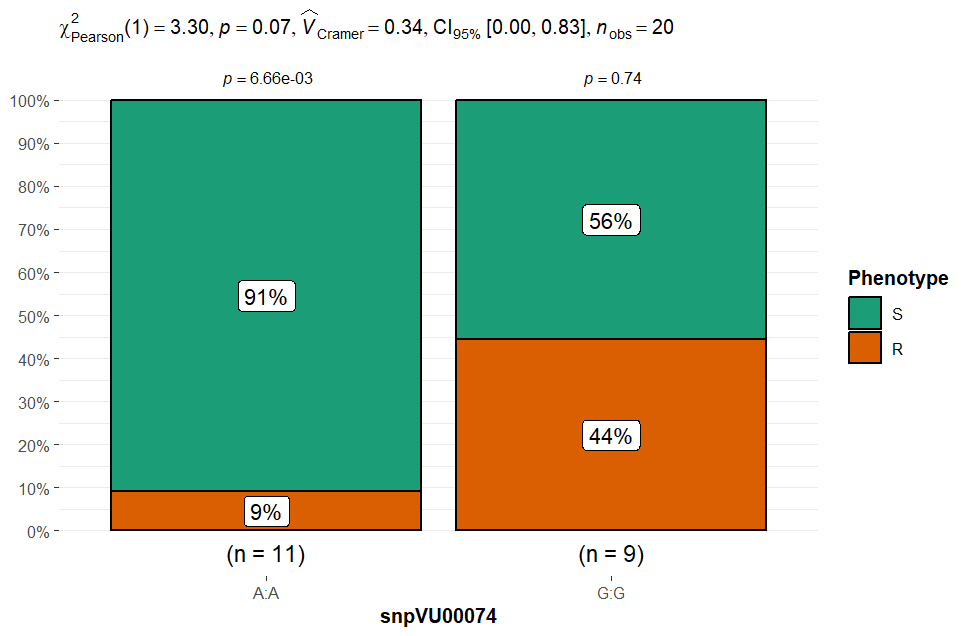

**Supplementary Figure 1C.** Chi-square test results for marker validation presented as stacked bar charts depicting the grouping of phenotypic reactions to Striga by the alleles of one candidate marker on chromosome Vu10. The 20 cowpea lines are grouped based on their phenotypic reaction to Striga into resistant (R) and susceptible (S) classes and the stacked bar charts are plotted to reflect the percentage of lines in each phenotypic category that carry the two alleles of a SNP marker. Summary statistics are presented at the top of each stacked bar chart that include chi square test (χ^2^), probability (p) presented for each marker allele class and for overall test of chi square test of independence, Cramer’s correlation (V̂_Cramer_), 5% confidence interval (CI_95%_) and number of observations (n_obs_).

Chromosome Vu11

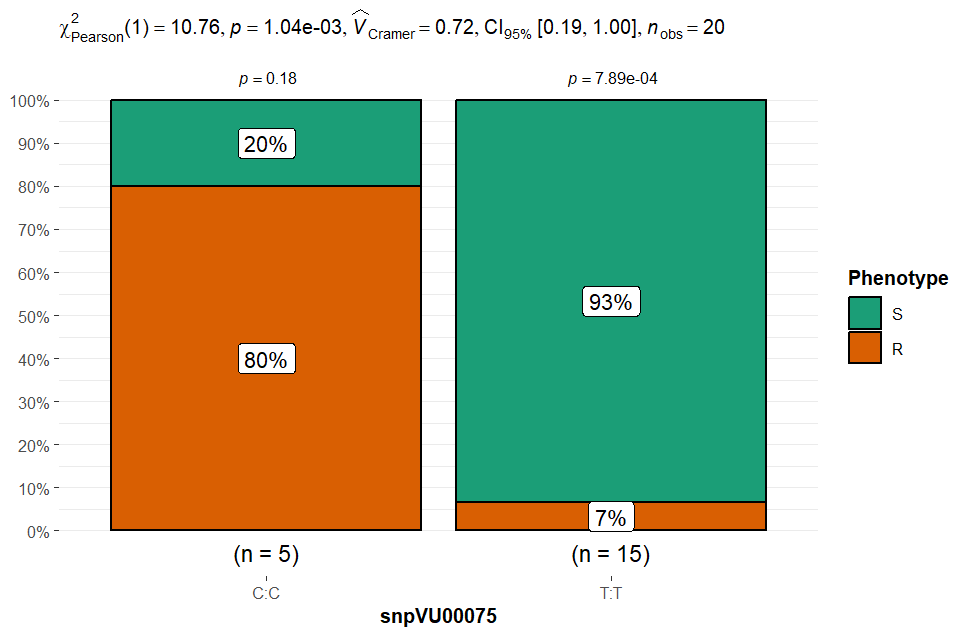

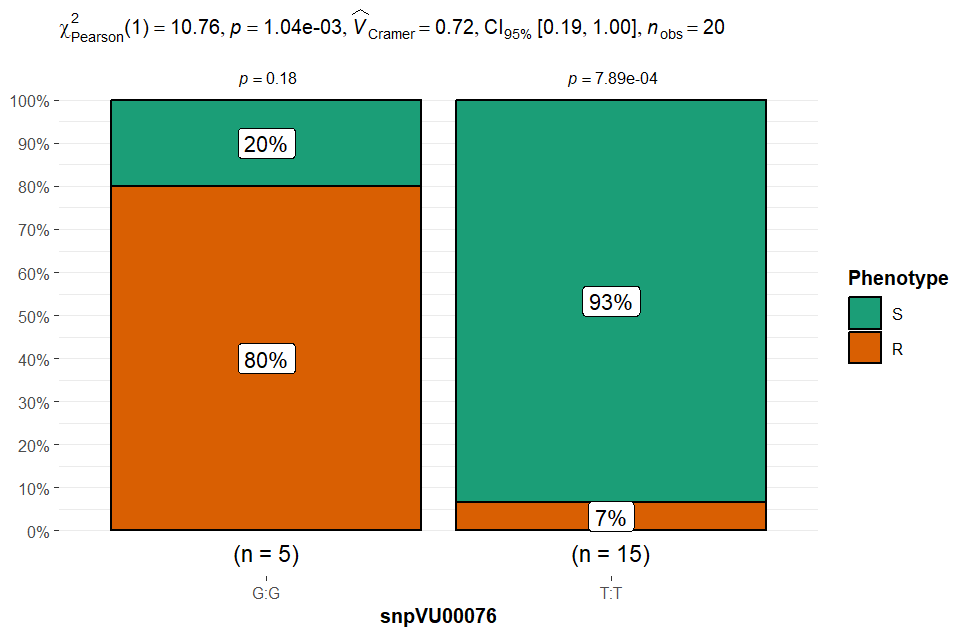

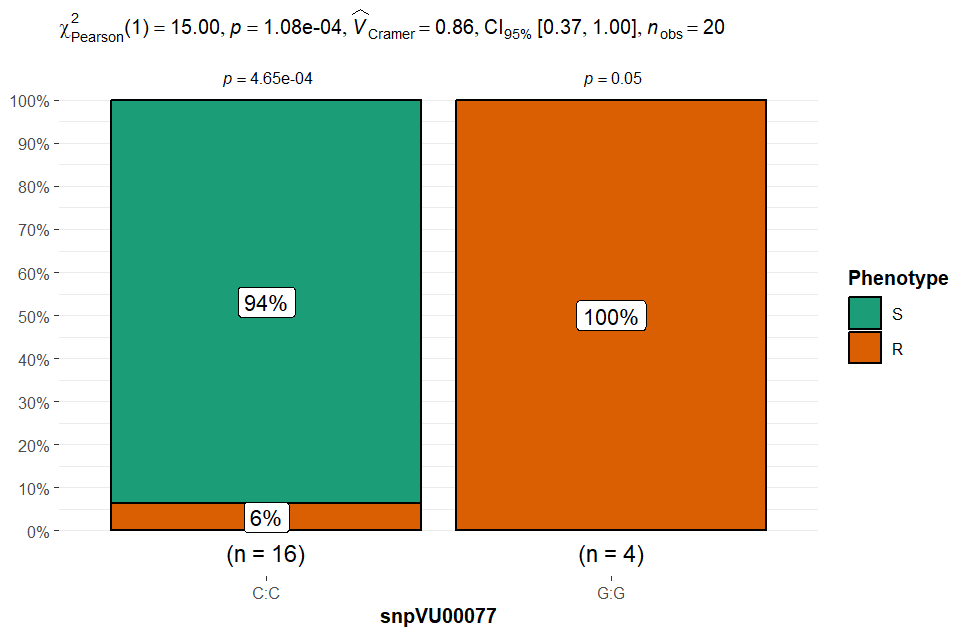

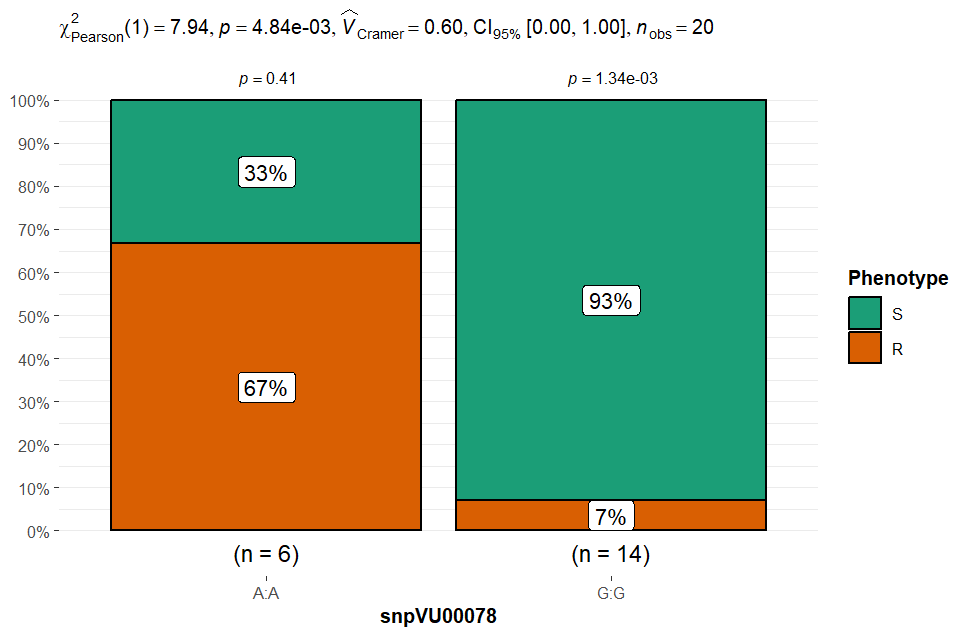

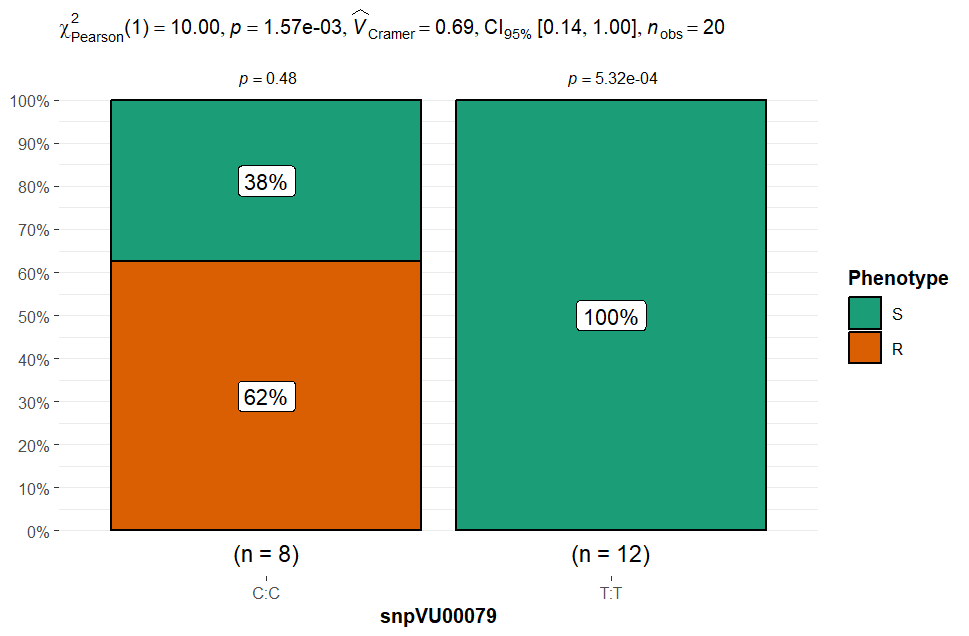

**Supplementary Figure 1D.** Chi-square test results for marker validation presented as stacked bar charts depicting the grouping of phenotypic reactions to Striga by the alleles of five proximal candidate markers on chromosome Vu11. The 20 cowpea lines are grouped based on their phenotypic reaction to striga into resistant (R) and susceptible (S) classes and the stacked bar charts are plotted to reflect the percentage of lines in each phenotypic category that carry the two alleles of a SNP marker. Summary statistics are presented at the top of each stacked bar chart that include chi square test (χ^2^), probability (p) presented for each marker allele class and for overall test of chi square test of independence, Cramer’s correlation (V̂_Cramer_), 5% confidence interval (CI_95%_) and number of observations (n_obs_).

### 1.2 Supplementary Figure 2

Chromosome Vu02

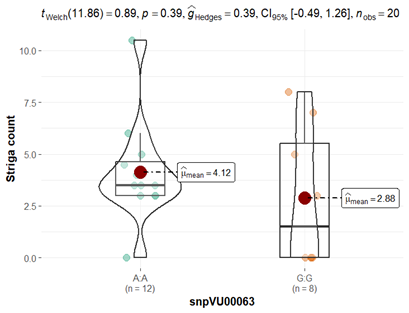

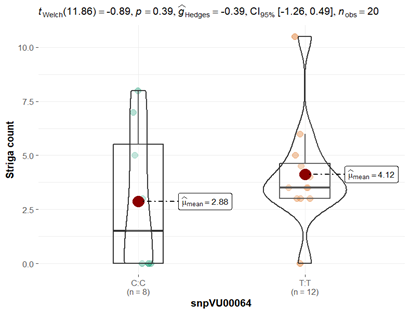

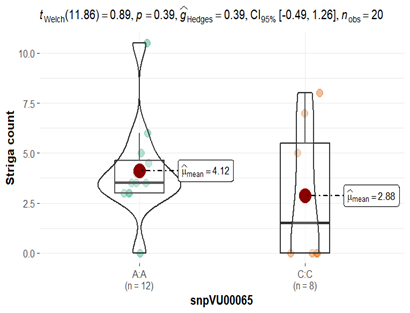

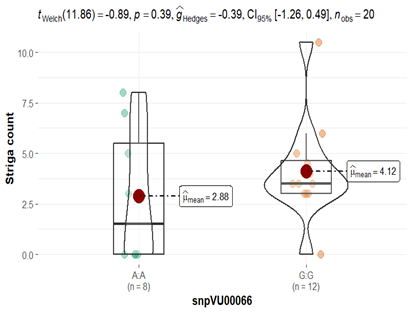

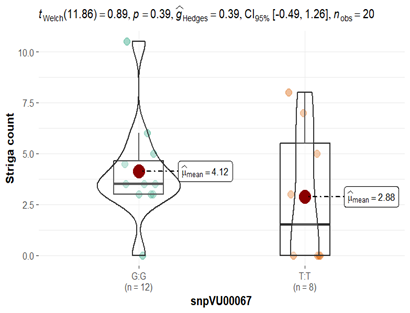

**Supplementary Figure 2A.** T-test results for marker validation presented as violin charts depicting the difference in mean Striga count between lines carrying the two alleles of each of the five proximal candidate markers on chromosome Vu02. The mean of each allelic group is indicated by a red dot and $\hat{\mu}$_mean_. Summary statistics are presented at the top of violin chart including t-statistical test (t_welch_), probability (p) for the t-test, effect size measured by Hedges’ g (ĝ_Hedges_), a 5% confidence interval (CI_95%_) and number of observations (n_obs_).

Chromosome Vu07

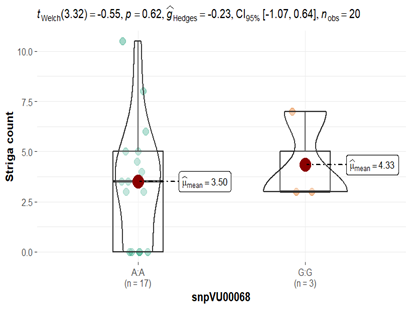

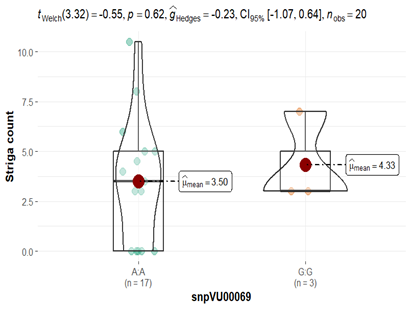

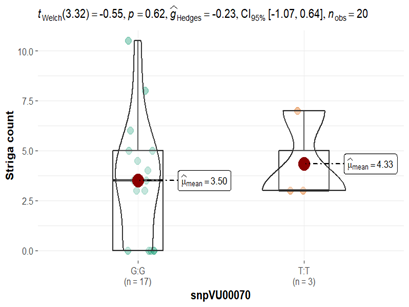

**Supplementary figure 2B.** T-test results for marker validation presented as violin charts depicting the difference in mean Striga count between lines carrying the two alleles of each of the three proximal candidate markers on chromosome Vu07. The mean of each allelic group is indicated by a red dot and $\hat{\mu}$_mean_. Summary statistics are presented at the top of violin chart including t-statistical test (t_welch_), probability (p) for the t-test, effect size measured by Hedges’ g (ĝ_Hedges_), a 5% confidence interval (CI_95%_) and number of observations (n_obs_).

Chromosome Vu10

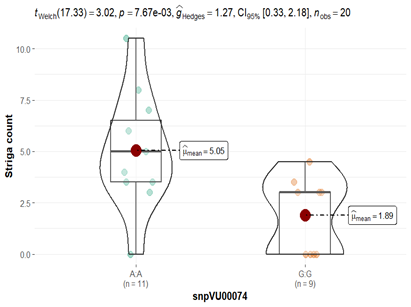

**Supplementary figure 2C.** T-test results for marker validation presented as violin charts depicting the difference in mean Striga count between lines carrying the two alleles one candidate marker on chromosome Vu10. The mean of each allelic group is indicated by a red dot and $\hat{\mu}$_mean_. Summary statistics are presented at the top of violin chart including t-statistical test (t_welch_), probability (p) for the t-test, effect size measured by Hedges’ g (ĝ_Hedges_), a 5% confidence interval (CI_95%_) and number of observations (n_obs_).

Chromosome Vu11

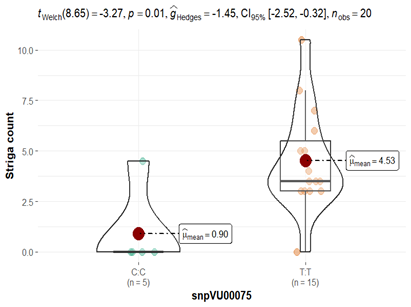

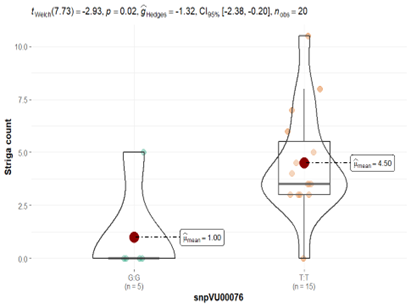

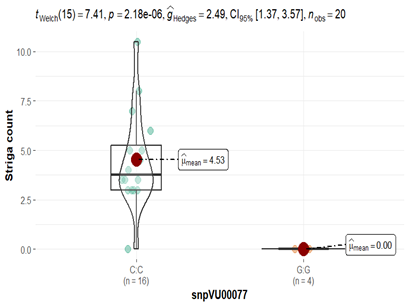

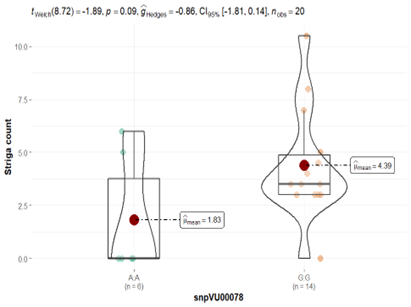

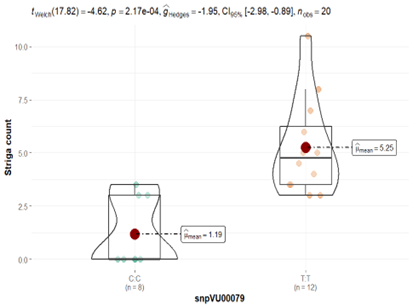

**Supplementary figure 2D.** T-test results for marker validation presented as violin charts depicting the difference in mean Striga count between lines carrying the two alleles of each of the five proximal candidate markers on chromosome Vu11. The mean of each allelic group is indicated by a red dot and $\hat{\mu}$_mean_. Summary statistics are presented at the top of violin chart including t-statistical test (t_welch_), probability (p) for the t-test, effect size measured by Hedges’ g (ĝ_Hedges_), a 5% confidence interval (CI_95%_) and number of observations (n_obs_).
